# Multiplicative frequency and angular speckle reduction in ultrasound imaging

**DOI:** 10.1101/2023.10.26.564267

**Authors:** Yilei Li, Noah Toyonaga, James Jiang, Alex Cable, Steven Chu

## Abstract

Speckle is the major artifact in ultrasound imaging, and it is well-known that speckle can be reduced by compounding (averaging) images taken either at different frequencies or from different angles. By averaging images of a phantom taken over a frequency range 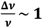 and a 90° span of angles, the combined speckle reduction is demonstrated to be ~ 9× compared to non-compounded images, while the reduction with frequency or angle averaging resulted in reductions of ~ 3× individually. The rf input to the transducer is altered to vary the sound frequency and the phantom is rotated with respect to the transducer to obtain different imaging angles. Numerical simulations of sound scattered by randomly distributed point scatterers showed quantitative agreement with the experiment. Using a commercial system, a 6× reduction in speckle is demonstrated imaging a human wrist. A robot arm is used to move the transducer along a circular path to acquire images at 9 angles separated by 10°. The commercial system does not allow direct control of the input to the transducer, so the broadband signal detected is Fourier filtered to obtain images at different frequencies with ~ 2× reduced frequency range. Images taken at different angles contain distortions from speed of sound variations and pressure induced by the probe. Two forms of non-rigid image registration are applied to correct for the distortions and create a higher resolution composite image. A design for achieving ~10× speckle reduction with essentially no loss in imaging speed is described.

## I. Introduction

Ultrasound imaging is an important medical diagnostics tool. It offers low cost, real-time imaging with no exposure to ionizing radiation. Refinements such as color Doppler, shear wave and contrast agent labeling offer valuable additional diagnostic information which complements x-ray, CT and MRI imaging modalities. However, ultrasound imaging is subject to significant speckle noise that severely degrades the effective resolution and limits its clinical use.

Speckle arises from the pulse-echo operation principle of ultrasound imaging. [1], [2] Each image, known as a B-scan, is formed with a sequence of A-scans, in which a sound pulse is transmitted, and scatters off the biological material in each voxel (imaging volume) as it propagates into the medium. The location of a corresponding voxel is determined from the time delay of the echo and the speed of sound. In a voxel, suppose there are (complex) scattering amplitudes 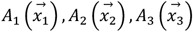. If these amplitudes interfere constructively or destructively, the scattered signal 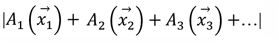 can be either more or less than the average of the scatterers, 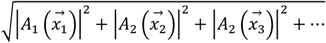. Even if each voxel has a high density of randomly distributed point scatters, the coherent interference of back-scattered signal may vary dramatically, thus producing significant speckle.

Physical methods to reduce speckle has been extensively investigated, where multiple images with varied speckle patterns are acquired by changing the imaging frequency (frequency compounding) or the angles of incidence (angle compounding). [3]–[20] The speckle noise can be reduced by a factor of up to 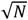 where *N* is the number of uncorrelated images taken at different frequencies or angles. It was found that a 2-4×reduction in speckle can be achieved in practice by either frequency [21]–[23] or angle averaging (where image averaging is sometimes referred to as compounding and spatial compounding is used interchangeably with angular compounding) [14], [15], [19], [24]. The maximal speckle reduction are limited physically by the bandwidth of the transducer [25] and the range of possible imaging angles. As a result, significant speckle noise is still present after the application of these methods. To the best of our knowledge, the improvement in speckle reduction when they are combined has not been examined.

There is also great interest in reducing speckle by image post-processing methods. [26]–[39] Some of the methods were applied to speckle noise reduction in synthetic aperture radar, which can be readily adapted to ultrasound imaging. In the adaptive filter approaches, an estimate of the level of speckle is computed in the vicinity of a target image location to determine the degree of smoothing [28]–[31], [34], [35], [40]–[42]. The anisotropic diffusion methods smooth an image in a process that mimics thermal diffusion [26], [27]. Various multiscale approaches utilizing wavelet or pyramid image decomposition have also been employed for speckle reduction.[43]–[54] By applying the post-processing methods on top of the physical speckle reduction methods such as frequency and angle compounding, further speckle reduction can be achieved.

Ultrasound transmission tomography reconstructs the attenuation coefficient and the speed of sound from the attenuation and the time of flight of sound pulses. [55]–[60] An advantage of transmission tomography is that speckle is absent since the secondary scattering amplitudes from each scatterer within the voxel add up in-phase in the forward direction. Moreover, the speed of sound map can be used to further correct for refraction in the reflection images. However, tomography requires the detector and transmitter to be located on the opposite sides of the object being imaged and as a result, its current primary application is in breast imaging.

In this work, we show that image compounding over a wide range of frequencies and angles reduces speckle in a statistically independent manner by factors of *X* ~ 3 and *Y ~* 3, and that the observed reduction agrees with a numerical simulation where the scattering medium is modeled as a collection of point scatterers. We also show that the speckle reduction of combined frequency and angle averaging is ~ 9, which is the product *XY*. We present a simple analytic argument to justify why the speckle reduction is consistent with the product when the interference patterns created by the point scatterers represent statistically independent speckle images. The combined frequency and angle compounding scheme is further implemented to image a human wrist using a robotic arm to take images at widely different angles. Non-rigid image matching algorithms are applied to correct for the mismatch between images taken from different angles. The multiplicative speckle reduction is also shown to pave the way for the achievement of more than 10×reduction in speckle in a practical imaging system.

## II. Physics of the multiplicative speckle reduction

In Fig. 1A and 1B, a simple argument is presented why speckle images at a fixed angle but different center frequencies must be inherently different than images taken at a different angle. In Fig. 1A, an ultrasound voxel is taken with a lateral resolution ∆*x* = *λ*/2*NA*~4λ and an axial resolution ∆*z* = ~2λ. For small numerical apertures, the acoustic wavefront within the voxel is approximately flat. Consider randomly distributed point scatterers within the voxel indicated by the circles that contribute to the coherent generation of the backscattered ultrasound signal. In Fig, 1B With a change in angle *δθ* > sin^%!^(*NA*), where *NA* is the numerical aperture of the focused sound wave, an additional random phase that varies over a range greater than 2π is acquired for each scatterer, rendering the speckle at the two angles independent, regardless of the frequency of sound, which gives rise to the multiplicative speckle reduction.

**Fig. 1.**
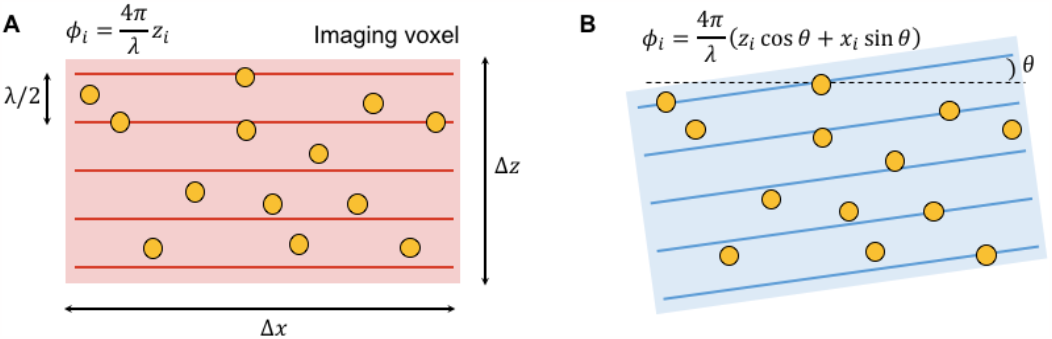
Coherent echo interference in a voxel. (**A**) An imaging voxel with randomly distributed scatterers (red rectangle). The axial and transverse dimensions of the voxel are ∆*z* and ∆*x*, respectively. In this example, the ultrasound voxel was assumed to have a lateral resolution ∆*x* = *λ*/2*NA*~4λ and an axial resolution 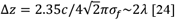. The titled lines indicates the snapshot in time of the wavefronts of a sound pulse, and the phases of the echo from each scattering point is a function of their axial position *z*_”_. The horizontal lines separated by *λ*/2 correspond to regions of the same constructive or destructive interference. By varying the ultrasound frequency *f*, the scattering planes that contribute to the constructive and destructive speckle interference are varied. If Δ*f* is defined as the range of frequencies from *f* symmetric spans Δ*f* = 2*f*, all the possible interference planes are sampled. An average of the voxel images is proportional to the total number of scatterers in the voxel. In Fig. 1B, (B) The imaging voxel (blue rectangle) with an angle of incidence rotated by *θ* relative to A.. The phases of the sound scattered off each scatterer are expressed in the coordinate system of A. Note that as *f* → 2*f*, the interference planes sampled in orientation B are independent of the planes sampled in orientation A when the relative angle *δθ* of the two orientations is greater than sin^−1^(*NA*), where *NA* is the numerical aperture of the focused sound wave.

We have previously shown that the maximal speckle reduction is achieved by using a number of evenly spaced Gaussian bands within a total bandwidth of ∆*f*. [25] The maximal speckle reduction achieved by frequency averaging (sometimes referred to as compounding) is approximately 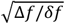, where *δf* is the full-width at half maximum of the Gaussian pulses. By averaging images taken over more than an octave of frequency range with *δf*~∆*f*/9, speckle noise can be reduced by up to ~3×. Similarly, in angle compounding, if the range of angles ∆θ allows for *N* different independent angles, the speckle reduction is 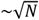 until the span of independent angles reaches 180°.

## III. Phantom experiment

Speckle reduction of combined frequency and angle compounding is quantified by imaging an agarose phantom with dispersed corn starch particles. The particles have typical sizes of 5 – 10 μm, much smaller than the ultrasound wavelength (λ ~ 200 – 700 μm) used here. A cylindrical hyperechoic region in the phantom has 3× corn starch concentration compared to the surrounding region.

### A. Phantom construction

The hypoechoic region is formed with 0.5 w/v (weight/volume) agarose mixed with 0.5 w/v corn starch. The agarose is dissolved in water by heating the mixture to boil in a microwave oven. Corn starch (0.5 w/v) is then mixed with the agarose solution and stirred for 30 minutes in a water bath at 50 ºC. Half of the solution is poured into a phantom container. An aluminum rod is placed in the container to be removed later to create a space for the formation of the hyperechoic region. Then the agarose solution with corn starch in the container is cooled down to room temperature to form gel. During this time, additional corn starch is added to the 0.5 w/v agarose solution so that the corn starch concentration reaches 1.5 w/v. The solution is placed in the water bath at 50 ºC and stirred to fully disperse the corn starch. As the phantom begins to gel, it is placed in a 4 ºC cold room for the gelation process to complete. After 3 hours, it is taken out of the cold room and warmed up to room temperature. The aluminum rod is then removed, and the 1.5 w/v agarose solution is poured into the phantom box to fill the space originally occupied by the rod. The phantom is left at room temperature to form gel and then placed in the 4 ºC cold room for the gelation of the hyperechoic region to complete.

### B. Image acquisition

A single element ultrasound transducer (Olympus C309-SU-F2.00IN-PTF) is used to image the phantom, where both the transducer and the phantom are submerged in water. The transducer has a focal distance of 5 cm and a diameter of 1.2 cm. A B-scan image is produced by a series of A-scans spatially separated by translating the transducer with a linear stage. A total of 100 lines are imaged with a spacing of 0.2 mm. (See Fig. S1A for a schematic of the experiment setup) The cross-section of the phantom corresponding to the imaging plane is illustrated in Fig. 2 (top panel). Ultrasound Gaussian pulses are generated by creating a digital waveform *A*_*dig*(_(*t*) of the desired pulse with an arbitrary waveform generator. The analogue electrical pulse is then amplified by a RF amplifier, which is gated in a time window Δ*t* = 10*σ*_*t*_ centered about the RF pulse so that the noise floor of the amplifier does not contribute to the detected signal *A*(*t*).

**Fig. 2.**
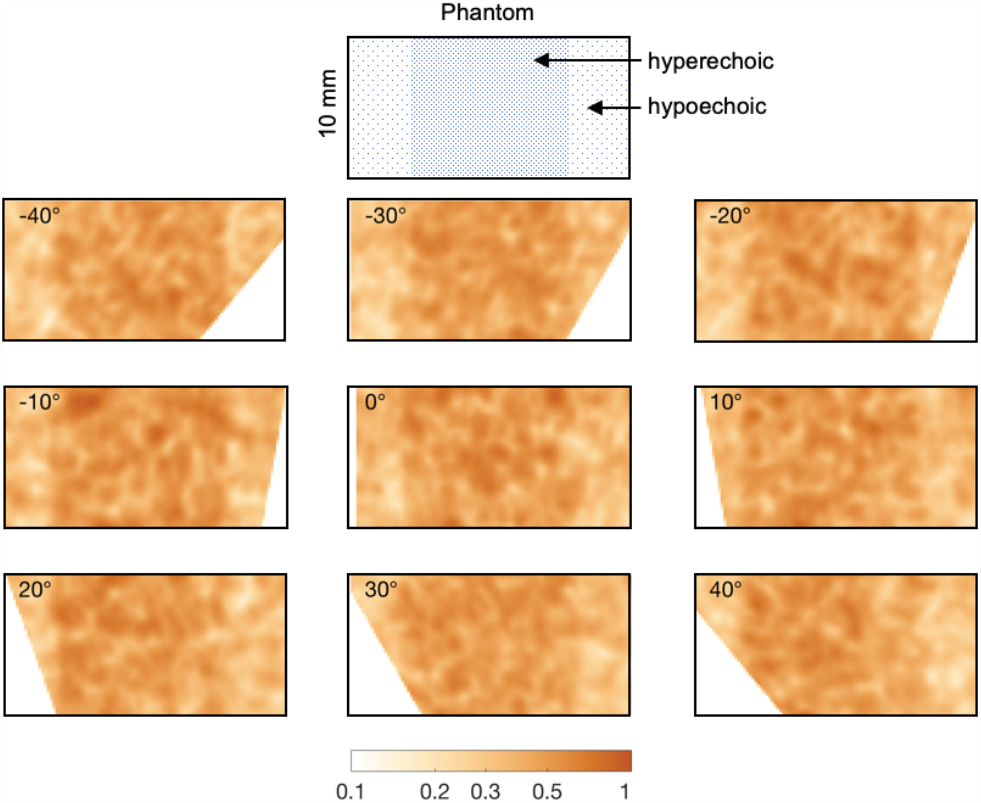
Frequency compounded phantom images from 9 angles. The images shown correspond to the same spatial region in the phantom. The center hyperechoic region has 3× higher corn starch concentration than the rest. The amplitude of the ultrasound is plotted on a false color scale shown at the bottom. A schematic of the phantom is shown above the experimental images.

To perform angle compounding, 9 images of the phantom are acquired by rotating the phantom to 9 different angles with 10° interval using a motorized rotation stage with its axis of rotation perpendicular to the imaging plane. Ultrasound Gaussian pulses centered at 9 different frequencies (2.4, 3.0, …, 7.2 MHz) are used to form the 9 images at each angle, so that the fractional spread in frequencies used was 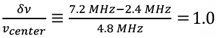. The pulses have a 1 *σ* time width *σ*_*t*_ and frequency width *σ*_*f*_ satisfying the minimum uncertainty 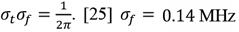 [25] *σ*_*f*_ =0.14 MHz is less than 1/4 of the center frequency spacing. As a result, the correlations in the speckle images obtained using the different frequency bands are insignificant.[3], [4], [23] The bandwidth gives an axial resolution (FWHM) of 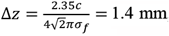, where *c* ≈ 1500 m/s is the speed of sound.[25] The diffraction limited lateral resolution is 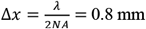, 1.2 mm, and 2.4 mm at 7.2 MHz, 4.8 MHz, and 2.4 MHz, respectively, where *NA* = 0.12 is the numerical aperture. The images are normalized to the average pixel intensity within the hyperechoic region. The frequency compounded images at each angle are then transformed to the same coordinate system by digital image rotation. (Fig. 2) The center of rotation is determined by aligning the corners of the hyperechoic region in the images. (Fig. S1B)

### C. Results

Images without compounding, with a 9-fold frequency compounding only, with a 9-fold angle compounding only, and with combined 9-fold frequency and 9-fold angle compounding are shown in Figs. 3A-D. Strikingly, the rectangular shape of the hyperechoic region stands out clearly with the multiplicative speckle reduction. Speckle noise in the hyperechoic region is evaluated using the dimensionless quantity *µ*/*σ*, where *µ* and *σ* are the mean and standard deviation of the ultrasound amplitude, respectively. In a situation of randomly distributed and densely packed point scatterers, our experiment yields µ/*σ* of approximately 1.9 for non-compounded images, in agreement with our numerical simulations and previous studies on speckle statistics[1], [2]. The reduction in speckle noise will be measured with respect to this value throughout this paper. The increase in *µ*/*σ* (and hence the reduction in speckle) in the combined compounding scheme is measured to be 8.8×, consistent with a multiplicative improvement (2.9 × 3.1 = 8.99) due to frequency and angle compounding of 2.9× and 3.1×, respectively.

**Fig. 3.**
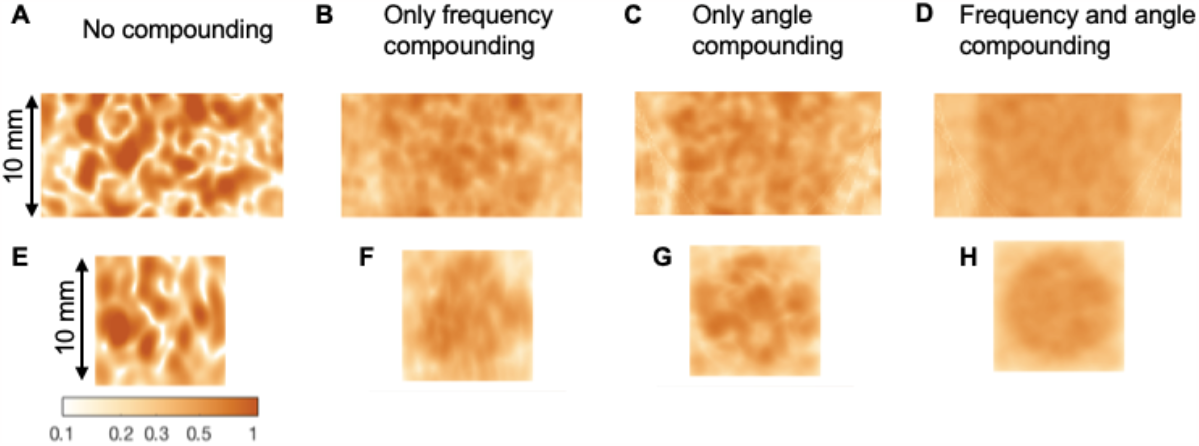
Experimental and simulated images without compounding, with either frequency or angle compounding, and with combined frequency and angle compounding. Images of the agarose phantom (A) at a single angle 0° and a single frequency 4.8 MHz. The axial (z) resolution is equal to the lateral (x) resolution. (B) Compounded image of averaging 9 frequencies centered at 2.4, 3.0, …, 7.2 MHz and a single angle 0°. (C) Compounded image of averaging 9 angles −40, −30, …, 40° and at a single frequency band centered at 4.8 MHz. (D) Compounded image obtained from 9 frequencies and 9 angles (a total of 81 images). (E-H) The simulated speckle with the corresponding conditions in (A-D).

## IV. Numerical simulation

Numerical simulation is performed with a distribution of point scatterers in a 10 mm by 10 mm region where a subregion (hyperechoic) contains a 3× higher density (600/mm^2^) than the rest (hypoechoic). The spatial distribution of scatterers within either region is random. Beyond a density of about 50 particles per voxel (~ 50/mm^2^), further increase in scatterer density does not result in appreciable change in the simulated speckle reduction figures. A snapshot of the pressure field *p* of the sound pulse is modeled by a Gaussian profile

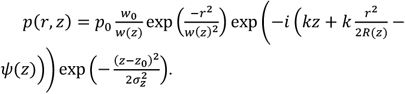

Here *r* is the radial distance from the center axis of the beam; *z* is the axial distance from the focus of the beam; *i* is the imaginary unit; *k* = 2π/λ is the wave number; *p*_>_ is the pressure at the origin; 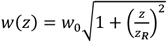 is the radius at which the field amplitude falls to 1/*e* of their axial values *(i*.*e*., where the intensity falls to 1/*e*^2^ of their axial values) and 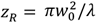 is known as the Rayleigh range; *w*_*0*_ = *w*(0) is the waist radius; 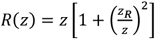 is the radius of curvature of the beam’s wave fronts at 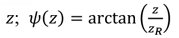 is the Gouy phase; *σ*_2_ = *cσ*_*t*_ is the pulse length. The numerical aperture of the Gaussian beam and its frequency bandwidth are chosen to match the experiment. The scatterers are assigned equal scattering cross-section and the scattered wave from individual scatterer is assumed to be a spherical wave. The received sound magnitude is calculated by summing the contribution from each scatterer and the spatial variation of speckle is obtained by scanning the origin of the Gaussian pulse profile over the simulated region. The calculation is performed using MATLAB.

The simulation results (Figs. 3E-H) show an improvement in image quality in accordance with the experiment. Quantitatively, the *µ*/*σ* increase of the combined compounding is 9.0×, whereas frequency or angle compounding alone increases *µ*/*σ* by 3.2× and 3.3×, respectively. The simulation results confirms the experimental finding that frequency and angle speckle reduction are multiplicative when combined.

## V. Recording images using a robot arm

Demonstration of the fundamental principle of multiplicative speckle reduction by combining frequency and angle compounding is concluded at this point, the focus of the rest of this paper is shifted towards the application of the scheme to image a human wrist. Using a commercial ultrasound system, images of a human wrist at specific angles are acquired by manipulating an ultrasound probe (MS250, VisualSonics, Inc.) with a robot arm (Meca500, Mecademic, Inc.) that has a spatial accuracy of 0.1 mm. The ultrasound probe is connected to an ultrasound imaging system (Vevo 2100, VisualSonics, Inc.). The robot positioned the probe at 9 radial positions separated by 10° around the circumference of the wrist. The acquired images are rotated digitally to account for the variation in imaging angle (Fig. 4). The ultrasound imaging system has a lateral resolution of ~ 0.2 mm and an axial resolution of ~0.1 mm. A layer of ultrasound gel of 1-2 mm thickness is applied to fill up the space between the probe and the wrist for acoustic coupling.

**Fig. 4.**
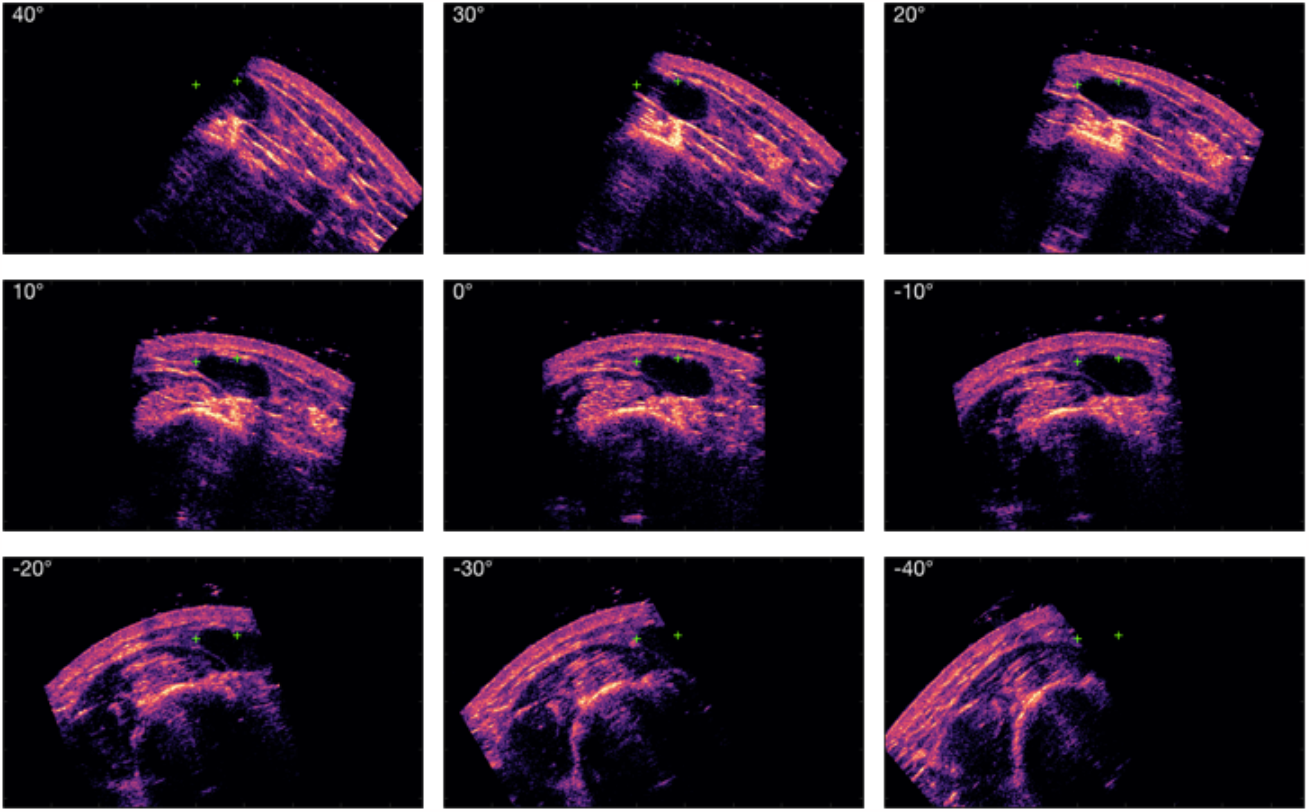
Frequency compounded images of the wrist (*σ*_*f*_ = 2.5 MHz). A common center of rotation for the set of images is determined by using the known angles from the robot so that the images of the blood vessel (the dark “oval-like” shape) overlap as much as possible. After accounting for the rotations, fiducial locations such as the green crosses in the sub-figures can be inserted in the images. The dimensions of the sub-figures are 43.5 mm ×26.0 mm.

In much of the previous work on angular compounding, the probe was held at a fixed position, and different portions of the transducer array were used to produce images from different angles [13], [14], [24]. For a phased array of total aperture length *L*, the resolution at a given depth *z* is approximately proportional to *L*/*z*. However, if the aperture is broken into *N* portions, the spatial aperture of each view is decreased by *N* and the resolution becomes (*L*/*N*)/*z, N* times worse than using the full aperture. Previously, feasibility of synthetic aperture imaging was examined by moving the transducer while precisely tracking the location and orientation of the transducer. [61]In addition, 3D ultrasound has been performed by combining 2D ultrasound images. [62], [63]

The VisualSonics imaging system does not allow independent control of the frequency spectrum of the ultrasound pulses. Frequency compounding is performed by Fourier filtering the back-scattered sound wave *A*(*t*) of each A-scan. *A*(*t*) is then Fourier transformed into the frequency domain *A*(*t*) ⟶ *F*(*f*), and digital Gaussian filters are applied to *F*(*f*) to generate *F*_*f*(*i*)_(*f*), where *f*(*i*) are the center frequencies of the spectrally filtered signals. The set of functions *F*_*f*(*i*)_(*f*) are then transformed back into the time domain to obtain *A*_*f*(*i*)_(*t*). The amplitude envelope of the radio frequency waveform is extracted using Hilbert transform.[64] In this way, 6 ultrasound images are produced, ranging from 6.0 MHz to 16.0 MHz, with spacings of ∆*f* = 2.0 MHz and bandwidths of *σ*_5_ = 2.5 MHz, with a corresponding spread in frequencies 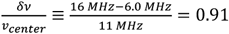. The reduced frequency range and the broader spectrum sound pulse used result in less speckle reduction compared to the data shown in Fig. 1. The subset of images at 6 different frequencies from each of the 9 angles are compounded to yield a set of 9 frequency compounded images.

## VI. Image distortions

Ultrasound images are distorted by a variety of factors including pressure from the ultrasound probe; refraction due to variations in the speed of sound in imaged media (Fig. 5). While not significant in this work, image distortions can potentially be induced by heartbeat, breathing, and other time-dependent variations in the tissue.

**Fig. 5.**
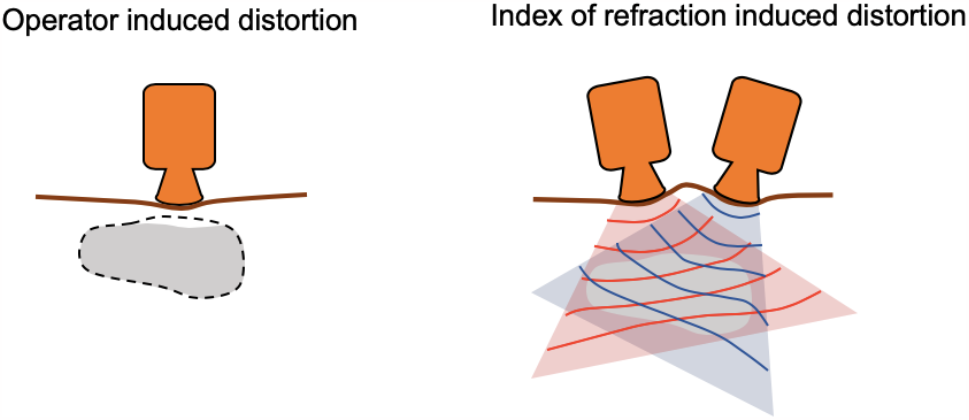
Factors that contribute to image distortions. Operator induce image distortion through the contact force of the probe with the patient. The dashed line shows the original location of the region marked in gray. The varying index of refraction in the imaging volume gives rise to weak lensing of the sound wave. The red and blue curves illustrate the wave front of a sound wave when the probe is located at two different orientations.

Image distortions reduce the effective resolution of the angle compounded image because features will not be correctly aligned across the set of images. This misalignment appears as blurring in the compounded image. By correcting these variations in structure, maximum resolution in the compounded image is maintained.

### A. Operator-induced variation

The operator can cause non-rigid geometric changes in the imaged volume during image acquisition. This can be a result of the pressure used to ensure good contact between the subject and the ultrasound probe. As the probe is moved to different locations, pressure exerted by the probe will distort the imaged volume into different configurations. In our experiment, an example of the distortion induced by the pressure of the ultrasound probe is shown in Fig. 6C. A comparison of two images taken 10° apart show mismatch after rigid image registration, most evident near the skin of the wrist indicated by the yellow arrow.

**Fig. 6.**
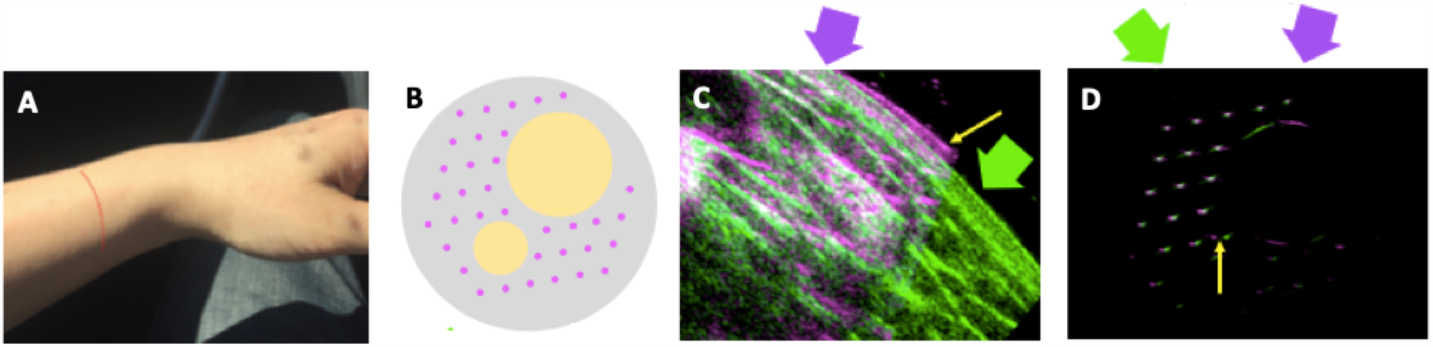
Observation of nonrigid distortions. (A) A photo of the imaged wrist. The red line indicates the location where the ultrasound images were acquired. (B) A schematic of the cross section of a 4 cm diameter phantom used to illustrate the distortion effects of sound refraction. The grey region is agarose and the yellow regions are filled with vegetable oil. The pink dots indicate the locations of 150 µ*m* diameter fishing lines. (C) Two scans (magenta and green, overlaid) after rigid image matching. Deformation near the skin surface (yellow arrow) is mostly due to pressure from the probe. (D) Overlaid images of the phantom taken from two different angles as indicated by the magenta and green arrows. The apparent location of a fishing line shifts due to refraction (yellow arrow). In (C) and (D), the false color images have dynamic ranges of 30 dB.

### B. Speed of sound variation

An ultrasound beam passing through a material with inhomogeneous speed of sound will be weakly lensed as it propagates through the material, analogous to looking through slightly “wavy” glass. This lensing effect changes depending on the imaging angle as the beam propagates through a different set of structures (and the corresponding indices of refraction).

In order to examine the effect of refraction, a cylindrical phantom (Fig. 6B) with regions of different speed of sound was constructed. The main body of the phantom is made of 1% w/v agarose water gel, where the speed of sound is approximately equal to the speed of sound in water. Two hollow cylindrical regions in the phantom are filled with vegetable oil by enclosing the phantom in a condom filled with vegetable oil. The speed of sound in vegetable oil is around 5-10% lower than that of water [65], comparable to the difference in the speed of sound between fatty and non-fatty soft tissues [66]. A hexagonal array of fishing lines (0.2 mm diameter) is stretched parallel to the phantom axis acting as fiducial markers. Images are taken perpendicular to the cylinder axis, and the lines appear as isolated point scatterers.

As the direction of the incident ultrasound waves is varied, the apparent locations of the fishing lines shift when imaged through the vegetable oil region. (Fig. 6D) In this test, mechanical distortion effects are minimized by submersing the phantom in water so that the ultrasound transducer is not in contact with the phantom. No displacement of the lines in the foreground is observed and as a result the mismatch between the images after rigid matching is caused purely by sound refraction.

## VII. Non-rigid image registration algorithms

Rigid registration (i.e. using exclusively translations and rotations) is insufficient in aligning the images taken from different angles accurately to within a voxel due to the distortion sources discussed in the previous section (Fig. S2). In order to recover the full inherent precision of ultrasound imaging, non-rigid image registration between pairs of images is used.

Each pair of images are initially aligned using rigid translation and rotation. One image remains fixed and is denoted as the stationary image *S*. The other image, to be distorted, is denoted as the moving image *M*. In this discussion, we define an “image” as a function that maps the spatial coordinates (*x, y*) to a set of image color values. In the case of a grey-scale ultrasound image, the image *S* is the mapping *S*(*x, y*) → *s*. The non-rigid image matching algorithm finds a transform *T* that operates on spatial coordinates of image *M* such that the resulting deformed image *M*′ matches with the stationary image *S* as closely as possible. (Mathematical descriptions of image and image transform are provided in Supplemental materials Sect. 2.1.)

Our non-rigid matching process starts with the image originally acquired at −40° as the stationary image *S*, while the image acquired at −30° is taken as the moving image *M*_−30°_. A non-rigid image algorithm is used to transform *M*_−30°_ → *M*_−30°_′. Next, the average of *S* and *M*_−30°_′ is set as the new stationary image *S*, and the image acquired at −20° becomes the new moving image *M*_−20°_ → *M*_−20°_′. The third *S* is the weighted average of *S* and *M*_−20°_′. This matching process continues until all images taken at different angles are added to the compounded image.

Two non-rigid image algorithms, the “demons” algorithm (so named in reference to Maxwell’s demons) [67] and stochastic gradient decent [68] are applied in this work (Table I). Both algorithms operate iteratively but differ in their ways to compute the updates to the image distortion *T*, which is described by a set of deformation vectors ***a***_*i*_ that gives the displacement of each pixel from the original image to the deformed image. The demons algorithm is based on local linear extrapolation of the intensity values while the gradient descent algorithm optimizes a global goodness-of-registration metric (in this experiment we use the mean squared error of intensity between images).

Both registration methods use “regularization” to constrain the solution to ensure robust, physically realistic solutions and prevent overfitting. If the parameters are under-constrained, an algorithm generates artificial registration maps which do not correspond to physical distortions and which are more sensitive to initial conditions. Over-constraining is equally problematic as it reduces the convergence speed of the algorithm and limits the space of attainable solutions and as a consequence the images are not matched fully. Examples of under-constrained and over-constrained regularization parameters are shown in Fig. S10. Further details about the image matching algorithms can be found in Sect. 2 of the Supplemental Material.

Fig. 7 shows the deformation vectors plotted on top of the images acquired from adjacent angles, where non-rigid deformation is applied to only one images in each pair. It can be seen that the direction and magnitude of the deformation vectors correspond to the mismatch of the features in the adjacent angle images, demonstrating the effectiveness of the non-rigid registration algorithm.

**Fig. 7.**
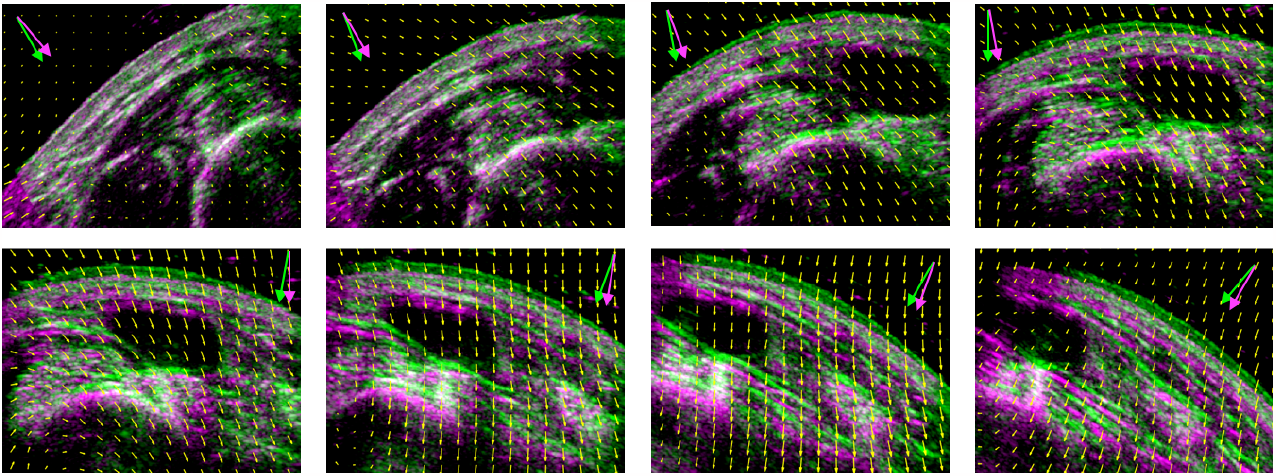
Adjacent angle images of a human wrist and the displacement vectors. The green and magenta arrows denote the imaging angles taken for each pair of images. The magenta images are corrected using the demons non-rigid image matching algorithm and the green images are the images taken at adjacent angles and aligned using only rigid translation and rotation. A subset of the displacement vectors are plotted on top of the images.

Figs. 8A and 8B compare the uncompounded images to the images with 6-frequency and 9-angle compounding using the demons algorithm. Figs. 8D and 8E show the comparison in a different region. Since the images taken from different angles do not overlap completely, the number of angles used in compounding varies spatially (Figs. 8C and 8F). The amount of speckle reduction expected from compounding 6 frequencies with *σ* = 2.5 MHz is ~1.3× [25] and that from compounding 9 angles is ~3.0× (assuming uncorrelated speckle in the images taken from different angles), giving a total reduction of ~3.9×. In this instance, the achieved speckle reduction from frequency compounding is significantly limited by the data acquisition bandwidth in the commercial ultrasound system. By reducing the 1*σ* pulse bandwidth to 1.2 MHz and hence relaxing the axial resolution to be comparable to the lateral resolution, the total speckle reduction is improved to 5.6×. We point out that the estimated speckle reduction in the wrist images are not directly obtained from the images. Fig. S4 shows the compounded images for different bandwidths and axial resolutions.

**Fig. 8.**
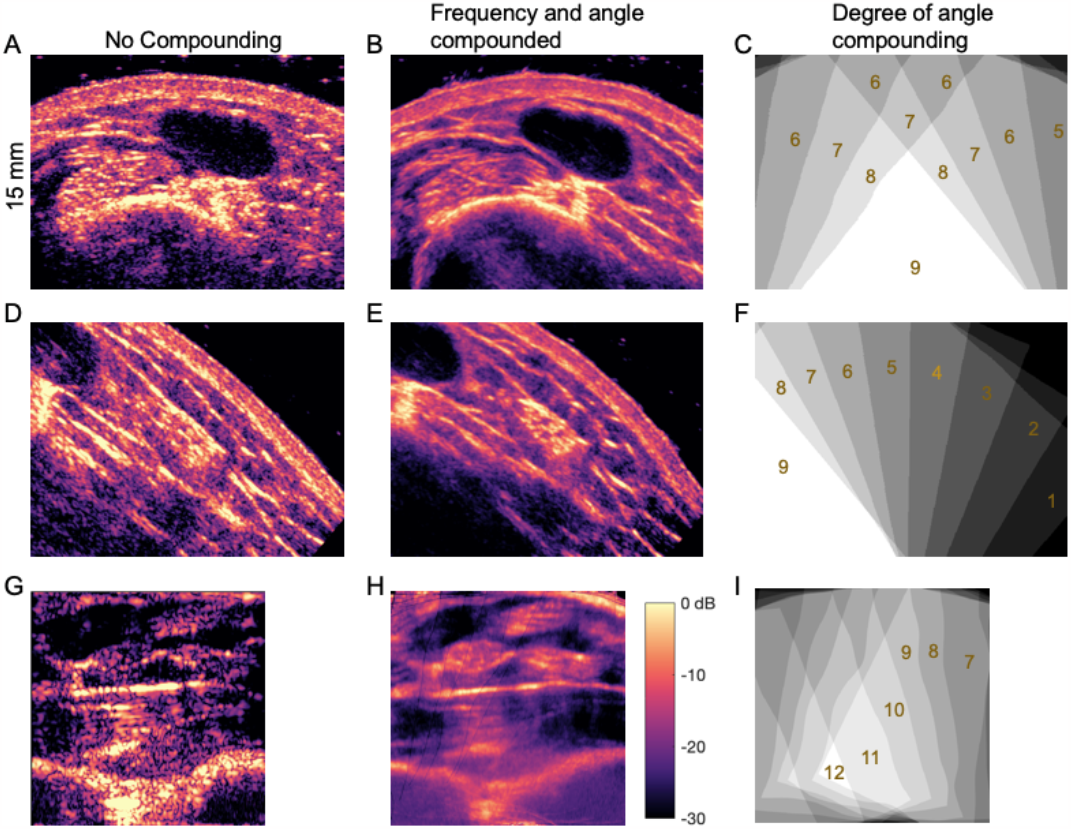
Imaging of human wrist. (A, D) False color images of a human wrist taken at different locations with a single angle and frequency. (B, E) Demons algorithm images compounded with 6 frequencies and 9 angles (a total of 54 images). (C, F) The spatial maps of the number of angles used for compounding corresponding to the ultrasound images in the same rows. For images A-F, the robot arm was used to acquire images at different angles and demons algorithm was used for image registration. Images G-H were acquired with a hand-held probe and gradient descent algorithm was used. (G) Image at a single angle and frequency. (H) The compounded image. Because of the flatter wrist geometry on the palm-side of the wrist, the overlap (I) of images is decreased. In Sect. VII Discussions, a practical engineering solution is proposed, which optimizes the overlap of the images using auto-beam steering over large variations in the imaging angle.

Both the demons and the gradient descent algorithms improved the contrast and resolution of the final compounded image by correcting for the probe-induced distortion and ultrasound refraction effects (Fig. S11). However, it was found that either algorithms was only able to partially compensate for the severe distortions due to the sudden index of refraction differences in the phantom image shown in Fig. 6B. The large oil-filled volume acts as a strong lens which shifts the apparent position of the point scatterers (Fig. 6D) when imaged in the magenta direction.

Compound imaging was also performed without the assistance of a robot arm. Here the ultrasound probe was hand-held by an operator and scanned along the palm side of the wrist. Without precise control of the position of the probe, the relative positions and orientations were found by maximizing the cross-correlations between the images, and the non-rigid image compounding was performed using the gradient descent algorithm. Figs. 8G and 8H compare the uncompounded and the frequency and angle compounded images. Fig. 8I shows the corresponding degree of angle compounding. In this data set, the speckle reduction from angle is smaller than the square root of the number of angles since the relative angles between the frames are generally smaller than the condition for independent speckle: sin *θ* > *NA*. Additionally, the resolution was limited by imperfections in the registration, which is significantly more complex due to the variations in pressure used by the operator when scanning by hand. The angle-compounded image given in Fig. 8H demonstrates that a hand-held scanner provides sufficient stability to create an improved angle compounded image. However, in most scanning applications, the topology of surface does not allow the recording of a number of overlapping images.

## VIII. Discussion

The maximum speckle reduction is achieved in this work by using Fourier transform limited pulses and the precise movement of a robotic arm. However, it is likely that comparable image improvements can be obtained with a single, broadband ultrasound pulse and with a hand-held device. For example, we will next extend the Fourier filtering method described in the VisualSonics imaging section of this paper to create a set of independent, self-normalized frequency images with a single ultrasound pulse. Dedicated fast Fourier transform integrated circuits can be used to process the reflected signal so that no sacrifice in imaging speed is expected due to frequency compounding. It should also be possible to dispense with a precisely controlled robotic arm to guide the ultrasound transducer. Images at significantly different incident angles of a fairly flat region such as the human abdomen can also be acquired without large tissue distortions.

Assuming the generation and detection of a wide bandwidth ultrasound pulses with a *Q*_eFF_ = 1.5 (See Sect. VIII. Discussion) that contains ~ 10 independent frequency bands and 10 image angles, the maximally achievable speckle reduction is expected to be ~10×. In this estimation, an aspect ratio of ∆*z*: ∆*x* = 1: 2 is assumed at the center of the spectrum. Note that the resulting angle compounded image may not converge on the correct image if there were distortions due to contact with the ultrasound head or refraction. However, in many medical imaging applications, that absolute size, shape or location of a feature is less important than maximizing the ability to resolve the smallest features that could be used by the clinician to identify anomalies such as the presence of the tumor or small tear in tissue.

### A. Frequency-compounding with spectrally independent and normalized ultrasound images taken with a single pulse

All ultrasound transducers have limited frequency response, and even broad band transducers have *Q*-factors *Q* ≡ *Δf* /*f* < 1.5. The finite *Q* of the transducer can be compensated for by digitally generating a short RF pulse with a frequency spectrum that has enhanced amplitudes in the portions of the spectrum where the frequency response is declining. The desired RF pulse is created by converting the digital waveform that compensates the natural transducer response to an analogue RF-pulse with a digital-to-analog converter. The generation of the RF-pulse can be considered to be part of the “Send + Receive Hardware” of the ultrasound machine in Fig. S12. Fig. 9A shows the natural response of the transducer measured by reflecting the sound pulse off an aluminum block placed at the focal point of the transducer. The flat-top pulse is compensated in the spectral range from 2.5 MHz to 7.0 MHz and a modified flat-top pulse is generated with a compensated rf pulse that is smoothed by a Gaussian with σ_*f*_ = 0.4 MHz. The resulting ultrasound pulse waveforms are shown in Fig. 9B.

**Fig. 9.**
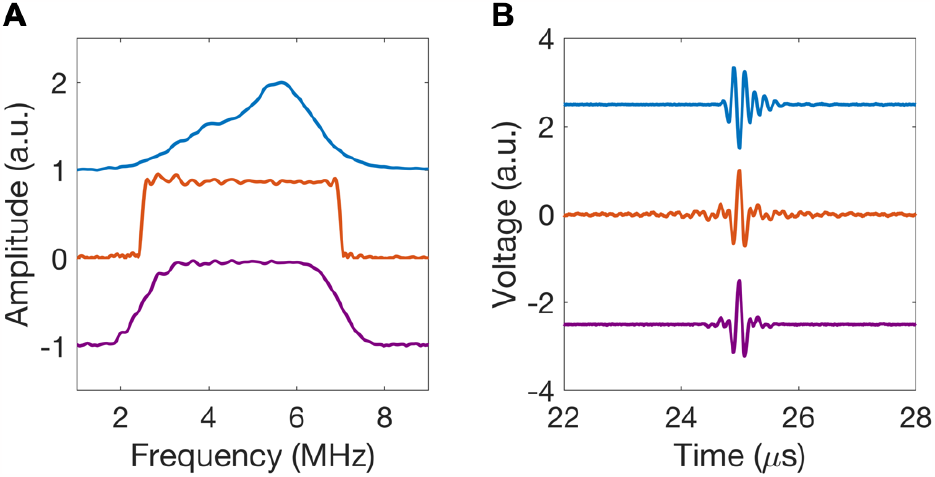
Generation of flat-topped ultrasound pulses. A. The natural response of the transducer (blue), the flat-top spectrum of a spectrally compensated sound pulse (red), and the flat-top spectrum of a spectrally compensated sound pulse where the rf excitation pulse is smoothed with a Gaussian function with *σ*_*f*_ = 0.4 MHz (purple). B. The time domain waveforms corresponding to the spectra presented in A.

Different frequency images are created by Fourier transforming the full-time record of the flat-topped broadband ultrasound echo signal *A*(*t*) into the frequency space: *A*(*t*) ⟶ *F*(*v*). The Fourier-transformed signal is then divided into a set of narrower-band digitally filtered signals centered at different frequencies *F*_*f*(*i*)_(*v*), and each filtered spectrum is Fourier transformed back into the time domain: *F*_*f*(*i*)_(*v*) ⟶ *A*_*f*(*i*)_(*t*) to create images centered at different frequencies to form images from a series of digitally filtered spectra. The separate images are compounded to form the reduced-speckle image.

### B. Forming angle compounded images with auto-steering

In order to minimize distortions due to the pressure of the ultrasound probe, the surface of the probe has to conform to the surface geometry of the patient. For example, in the case of imaging the palm-side of the wrist in Fig. 8H, the overlap of images as shown in Fig. 8I were limited because the topology of the wrist does not allow rotation of the probe about a single axis.

In future work, an automated beam steering procedure will be implemented to capture multiple angles of a region of interest (ROI) while minimizing operator-induced distortion. Fig. 10 illustrates one method of automatically using a phased-array beam to steer the ultrasound array to capture images over a large range of angles. After the ROI is identified by the clinician with the blue colored B-scan, the operator will initiate the start of a compound scan by, for example, pressing a foot-pedal. Without beam steering, the green image is captured after a small lateral displacement of the ultrasound transducer (Fig. 10A). By following the contour of the abdomen, the overlap of the two images is compromised.

**Fig. 10.**
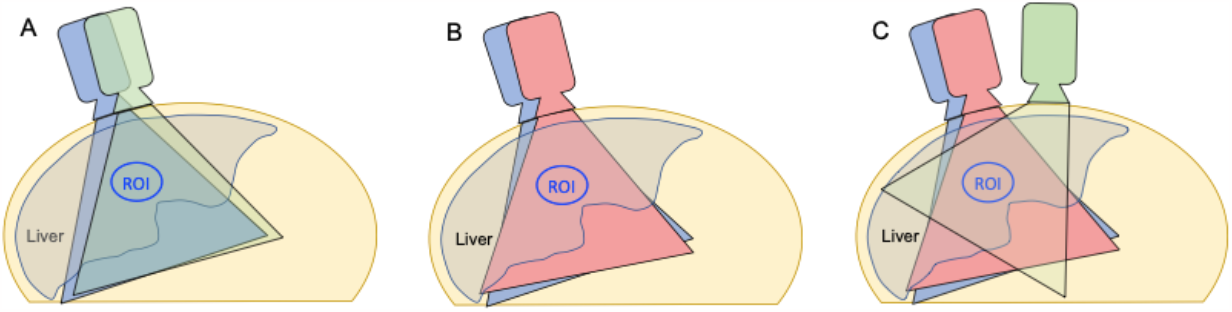
For a region of interest underneath a relatively flat surface such as a patient’s abdomen, images taken at significantly different angles would require excessive distortions. Instead, a phased array can be used to direct the ultrasound beam to image the ROI as the probe is moved across the patient’s abdomen. (**A**) Without beam steering, the image region will gradually move away from the ROI as the probe is moved. (**B**) Automatic beam steering corrects for the movement of the probe to direct the ultrasound beam to image the ROI. (**C**) Imaging of the same ROI is achieved over a large range of angles using automatic beam steering.

To optimize the overlap in the images taken from different angles, a phased array can be used to automatically steer the direction of the ultrasound B-scan to remain centered around the ROI. After an initial B-scan is selected by the clinician, the software cross-correlation of adjacent images shown in blue and green B-scans in Fig. 10A can be used to acquire maximally overlapped images as depicted in Fig. 10B. In this way, images taken over angles differing by ~90° can be acquired (Fig. 10C). Since large changes in angle is required to achieve significant speckle reduction in angle compounding, only a subset of the real-time images taken at angles θ_*i*_ are selected for compounding. Suppose the ultrasound machine is operating at a B-scan frame rate of 10 frames/second. A compound scan over 90 degrees with a threshold 10 degrees would need ~ 11 of the ~ 20 images taken over a span of 2 seconds.

It should be possible to achieve combined frequency and angle compounding imaging without substantially slowing down the data acquisition time used to manually scan a ROI. Currently, the image registration process takes a few seconds for each pair of images. Fast Fourier transform and graphic processing chips typically enables a parallelization factor of 100×or more, so frequency and angle compounding appear to be fast enough that the round-trip transit time of sound, instead of the speed of computation, limits the frame rate of the optimized system.

In a clinic application, the ultrasound operator will need to move a 2D imaging ultrasound probe along the direction of the 1D sensor array, or to rotate the probe within the 2D imaging plane. Failure of optimized overlap may be caused by moving the probe too quickly such that the adjacent frames are separated by more than the desired separation angle for compounding or when the probe is moved significantly away from the plane of imaging. A software routine running in the background will decide if image registration is successful or notify the operator to slow down the scan motion if the registration has failed. The process of acquiring a high-quality ultrasound compounded image can therefore guide the human operator without the aid of accurate probe positioning or measuring hardware (Fig. S12). In conclusion, minimal changes to the clinical practice of ultrasound imaging can result in substantially improved images.

## IX. Conclusion

In this work, we described the physical mechanism for the multiplicative speckle reduction by combining frequency and angle compounding. Experimentally, we demonstrate a **~**9×speckle reduction using a phantom of randomly distributed corn starch in agarose. A human wrist is imaged with ~6× reduction in speckle using a commercial system. The lower speckle reduction value in the latter case is due to the limited bandwidth of the commercial system, which lowers the speckle reduction with frequency. Image distortion mechanism including operator induced distortion, patient motion, and refractive index induced sound refraction are shown as sources of mismatch among images when performing angle compounding. Two non-rigid image matching algorithms are applied to correct for these distortions. We also describe a pathway towards achieving~10×reduction in speckle in an ultrasound imaging system with essentially no loss in imaging speed.

## Supporting information

http://www.stanford.edu/group/chugroup/

## Acknowledgment

We acknowledge the use of the VisualSonics Vevo2100 ultrasound scanner in the Small Animal Imaging Facilities at Stanford University.

## References

[1] C. B. Burckhardt, “Speckle in ultrasound B-mode scans,” IEEE Trans. Sonics Ultrason., vol. 25, no. 1, pp. 1–6, 1978.

[2] R. F. Wagner, S. W. Smith, J. M. Sandrik, and H. Lopez, “Statistics of Speckle in Ultrasound B-Scans,” IEEE Trans. Sonics Ultrason., vol. 30, no. 3, pp. 156–163, 1983.

[3] P. A. Magnin, O. T. von Ramm, and F. L. Thurstone, “Frequency compounding for speckle contrast reduction in phased array images,” Ultrason. Imaging, vol. 4, no. 3, pp. 267–281, 1982.

[4] H. E. Melton and P. A. Magnin, “A-mode speckle reduction with compound frequencies and compound bandwidths,” Ultrason. Imaging, vol. 6, no. 2, pp. 159–173, 1984.

[5] I. S. Song, C. H. Yoon, G. D. Kim, Y. Yoo, and J. H. Chang, “Adaptive frequency compounding for speckle reduction,” in IEEE International Ultrasonics Symposium, IUS, 2011.

[6] J. J. Dahl, D. A. Guenther, and G. E. Trahey, “Adaptive imaging and spatial compounding in the presence of aberration,” IEEE Trans. Ultrason. Ferroelectr. Freq. Control, 2005.

[7] J. Lee and J. H. Chang, “Dual-Element Intravascular Ultrasound Transducer for Tissue Harmonic Imaging and Frequency Compounding: Development and Imaging Performance Assessment,” IEEE Trans. Biomed. Eng., 2019.

[8] H. Tu, J. A. Zagzebski, A. L. Gerig, Q. Chen, E. L. Madsen, and T. J. Hall, “Optimization of angular and frequency compounding in ultrasonic attenuation estimations,” J. Acoust. Soc. Am., 2005.

[9] M. W. Urban, A. Alizad, and M. Fatemi, “Vibro-acoustography and multifrequency image compounding,” Ultrasonics, 2011.

[10] J. S. Ullom, M. Oelze, and J. R. Sanchez, “Ultrasound speckle reduction using coded excitation, frequency compounding,and postprocessing despeckling filters,” in Proceedings - IEEE Ultrasonics Symposium, 2010.

[11] Z. Zhang, L. Li, and H. Liu, “Ultrasonic elastography optimization algorithm based on coded excitation and spatial compounding,” Autom. Control Comput. Sci., 2017.

[12] G. Cincotti, G. Loi, and M. Pappalardo, “Frequency decomposition and compounding of ultrasound medical images with wavelet packets,” IEEE Trans. Med. Imaging, 2001.

[13] M. Berson, A. Roncin, and L. Pourcelot, “Compound scanning with an electrically steered beam,” Ultrason. Imaging, vol. 3, no. 3, pp. 303–308, 1981.

[14] G. E. Trahey, S. W. Smith, and O. T. von Ramm, “Speckle Pattern Correlation with Lateral Aperture Translation: Experimental Results and Implications for Spatial Compounding,” IEEE Trans. Ultrason. Ferroelectr. Freq. Control, vol. 33, no. 3, pp. 257–264, 1986.

[15] S. K. Jespersen, J. E. Wilhjelm, and H. Sillesen, “Multi-Angle Compound Imaging,” Ultrason. Imaging, vol. 20, no. 2, pp. 81–102, Apr. 1998.

[16] J. F. Krücker, C. R. Meyer, G. L. LeCarpentier, J. B. Fowlkes, and P. L. Carson, “3D spatial compounding of ultrasound images using image-based nonrigid registration,” Ultrasound Med. Biol., 2000.

[17] M. O’Donnell and S. D. Silverstein, “Optimum Displacement for Compound Image Generation in Medical Ultrasound,” IEEE Trans. Ultrason. Ferroelectr. Freq. Control, 1988.

[18] A. R. Groves and R. N. Rohling, “Two-dimensional spatial compounding with warping,” Ultrasound Med. Biol., 2004.

[19] G. E. Trahey, J. W. Allison, S. W. Smith, and O. T. von Ramm, “Speckle Reduction Achievable By Spatial Compounding And Frequency Compounding: Experimental Results And Implications For Target Detectability,” 1987, vol. 0768, no., p. 768.

[20] C. Yoon, G. D. Kim, Y. Yoo, T. K. Song, and J. H. Chang, “Frequency equalized compounding for effective speckle reduction in medical ultrasound imaging,” Biomed. Signal Process. Control, 2013.

[21] G. E. Trahey, J. W. Allison, S. W. Smith, and O. T. von Ramm, “A quantitative approach to speckle reduction via frequency compounding,” Ultrason. Imaging, vol. 8, no. 3, pp. 151–164, 1986.

[22] S. M. Gehlbach and F. G. Sommer, “Frequency Diversity Speckle Processing,” Ultrason. Imaging, vol. 9, no. 2, pp. 92–105, Apr. 1987.

[23] G. E. Trahey, J. W. Allison, S. W. Smith, and O. T. von Ramm, “Speckle Pattern Changes with Varying Acoustic Frequency: Experimental Measurement and Implications for Frequency Compounding,” in IEEE 1986 Ultrasonics Symposium, 1986, pp. 815–818.

[24] D. P. Shattuck and O. T. von Ramm, “Compound scanning with a phased array,” Ultrason. Imaging, vol. 4, no. 2, pp. 93–107, 1982.

[25] Y. Li, Y. Winetraub, O. Liba, A. De La Zerda, and S. Chu, “Optimization of the trade-off between speckle reduction and axial resolution in frequency compounding,” IEEE Trans. Med. Imaging, vol. 38, no. 1, 2019.

[26] Y. Yu and S. T. Acton, “Speckle reducing anisotropic diffusion,” IEEE Trans. Image Process., vol. 11, no. 11, pp. 1260–1270, 2002.

[27] K. Z. Abd-Elmoniem, A. B. M. Youssef, and Y. M. Kadah, “Realtime speckle reduction and coherence enhancement in ultrasound imaging via nonlinear anisotropic diffusion,” IEEE Trans. Biomed. Eng., vol. 49, no. 9, pp. 997–1014, 2002.

[28] P. C. Tay, S. T. Acton, and J. A. Hossack, “A stochastic approach to ultrasound despeckling,” in 2006 3rd IEEE International Symposium on Biomedical Imaging: From Nano to Macro - Proceedings, 2006.

[29] P. C. Tay, S. T. Acton, and J. A. Hossack, “Ultrasound despeckling using an adaptive window stochastic approach,” in Proceedings - International Conference on Image Processing, ICIP, 2006.

[30] D. T. Kuan, A. A. Sawchuk, T. C. Strand, and P. Chavel, “Adaptive Restoration of Images with Speckle,” IEEE Transactions on Acoustics, Speech, and Signal Processing. 1987.

[31] A. Lopes, R. Touzi, and E. Nezry, “Adaptive Speckle Filters and Scene Heterogeneity,” IEEE Trans. Geosci. Remote Sens., 1990.

[32] T. Loupas, W. N. McDicken, T. Anderson, and P. L. Allan, “Development of an advanced digital image processor for real-time speckle suppression in routine ultrasonic scanning,” Ultrasound Med. Biol., vol. 20, no. 3, pp. 239–249, Jan. 1994.

[33] K. Krissian, C. F. Westin, R. Kikinis, and K. G. Vosburgh, “Oriented speckle reducing anisotropic diffusion,” IEEE Trans. Image Process., 2007.

[34] J. Sen Lee, “Digital Image Enhancement and Noise Filtering by Use of Local Statistics,” IEEE Trans. Pattern Anal. Mach. Intell., 1980.

[35] V. S. Frost, J. A. Stiles, K. S. Shanmugan, and J. C. Holtzman, “A Model for Radar Images and Its Application to Adaptive Digital Filtering of Multiplicative Noise,” IEEE Trans. Pattern Anal. Mach. Intell., vol. PAMI-4, no. 2, pp. 157–166, 1982.

[36] P. C. Tay, C. D. Garson, S. T. Acton, and J. A. Hossack, “Ultrasound despeckling for contrast enhancement,” IEEE Trans. Image Process., 2010.

[37] M. A. Lediju, G. E. Trahey, B. C. Byram, and J. J. Dahl, “Short-lag spatial coherence of backscattered echoes: imaging characteristics,” IEEE Trans. Ultrason. Ferroelectr. Freq. Control, vol. 58, no. 7, pp. 1377–1388, 2011.

[38] Y. Chen, R. Yin, P. Flynn, and S. Broschat, “Aggressive region growing for speckle reduction in ultrasound images,” Pattern Recognit. Lett., 2003.

[39] H. C. Huang, J. Y. Chen, S. De Wang, and C. M. Chen, “Adaptive ultrasonic speckle reduction based on the slope-facet model,” Ultrasound Med. Biol., 2003.

[40] P. Coupé, P. Hellier, C. Kervrann, and C. Barillot, “Nonlocal means-based speckle filtering for ultrasound images,” IEEE Trans. Image Process., 2009.

[41] T. C. Aysal and K. E. Barner, “Rayleigh-maximum-likelihood filtering for speckle reduction of ultrasound images,” IEEE Trans. Med. Imaging, 2007.

[42] J. Jai Jaganath Babu and G. Florence Sudha, “Adaptive speckle reduction in ultrasound images using fuzzy logic on Coefficient of Variation,” Biomed. Signal Process. Control, 2016.

[43] F. Zhang, Y. M. Yoo, L. M. Koh, and Y. Kim, “Nonlinear diffusion in laplacian pyramid domain for ultrasonic speckle reduction,” IEEE Trans. Med. Imaging, 2007.

[44] Y. Yue, M. M. Croitoru, A. Bidani, J. B. Zwischenberger, and J. W. Clark, “Nonlinear multiscale wavelet diffusion for speckle suppression and edge enhancement in ultrasound images,” IEEE Trans. Med. Imaging, 2006.

[45] Z. Fan, M. Y. Yang, Z. Lichen, L. M. Koh, and Y. Kim, “Multiscale nonlinear diffusion and shock filter for ultrasound image enhancement,” in Proceedings of the IEEE Computer Society Conference on Computer Vision and Pattern Recognition, 2006.

[46] J. Kang and Y. Yoo, “New multiscale speckle suppression and edge enhancement with nonlinear diffusion and homomorphic filtering for medical ultrasound imaging,” in Medical Imaging 2014: Image Processing, 2014.

[47] S. Sudha, G. R. Suresh, and R. Sukanesh, “Speckle Noise Reduction in Ultrasound Images by Wavelet Thresholding based on Weighted Variance,” Int. J. Comput. Theory Eng., 2009.

[48] S. Gupta, L. Kaur, R. C. Chauhan, and S. C. Saxena, “A wavelet based statistical approach for speckle reduction in medical ultrasound images,” in IEEE Region 10 Annual International Conference, Proceedings/TENCON, 2003.

[49] Y. S. Kim and J. B. Ra, “Improvement of ultrasound image based on wavelet transform: speckle reduction and edge enhancement,” in Medical Imaging 2005: Image Processing, 2005.

[50] H. Rabbani, M. Vafadust, P. Abolmaesumi, and S. Gazor, “Speckle noise reduction of medical ultrasound images in complex wavelet domain using mixture priors,” IEEE Trans. Biomed. Eng., 2008.

[51] Xiaohui Hao, Shangkai Gao, and Xiaorong Gao, “A novel multiscale nonlinear thresholding method for ultrasonic speckle suppressing,” IEEE Trans. Med. Imaging, 1999.

[52] A. Achim, A. Bezerianos, and P. Tsakalides, “Novel Bayesian multiscale method for speckle removal in medical ultrasound images,” IEEE Trans. Med. Imaging, 2001.

[53] G. Andria, F. Attivissimo, A. M. L. Lanzolla, and M. Savino, “A suitable threshold for speckle reduction in ultrasound images,” IEEE Trans. Instrum. Meas., 2013.

[54] X. Zong, A. F. Laine, and E. A. Geiser, “Speckle Reduction and Contrast Enhancement of Echocardiograms via Multiscale Nonlinear Processing,” IEEE Trans. Med. Imaging, 1998.

[55] J. W. Wiskin, D. T. Borup, E. Iuanow, J. Klock, and M. W. Lenox, “3-D Nonlinear Acoustic Inverse Scattering: Algorithm and Quantitative Results,” IEEE Trans. Ultrason. Ferroelectr. Freq. Control, 2017.

[56] B. Malik, J. Klock, J. Wiskin, and M. Lenox, “Objective breast tissue image classification using Quantitative Transmission ultrasound tomography,” Sci. Rep., 2016.

[57] J. Wiskin, D. T. Borup, S. A. Johnson, and M. Berggren, “Nonlinear inverse scattering: High resolution quantitative breast tissue tomography,” J. Acoust. Soc. Am., 2012.

[58] M. André, J. Wiskin, D. Borup, S. Johnson, H. Ojeda-Fournier, and L. Olson, “Quantitative volumetric breast imaging with 3D inverse scatter computed tomography.,” Conf. Proc. IEEE Eng. Med. Biol. Soc., 2012.

[59] N. Duric, P. Littrup, L. Poulo, A. Babkin, R. Pevzner, E. Holsapple, O. Rama, and C. Glide, “Detection of breast cancer with ultrasound tomography: First results with the Computed Ultrasound Risk Evaluation (CURE) prototype,” Med. Phys., 2007.

[60] C. Li, N. Duric, P. Littrup, and L. Huang, “In vivo Breast Sound-Speed Imaging with Ultrasound Tomography,” Ultrasound Med. Biol., 2009.

[61] N. Bottenus, W. Long, H. K. Zhang, M. Jakovljevic, D. P. Bradway, E. M. Boctor, and G. E. Trahey, “Feasibility of Swept Synthetic Aperture Ultrasound Imaging,” IEEE Trans. Med. Imaging, 2016.

[62] D. H. Iversen, F. Lindseth, G. Unsgaard, H. Torp, and L. Lovstakken, “Improved quality of freehand 3-D ultrasound color flow imaging by multi-angle compounding,” 2015.

[63] C. D. Herickhoff, M. R. Morgan, J. S. Broder, and J. J. Dahl, “Low-cost Volumetric Ultrasound by Augmentation of 2D Systems: Design and Prototype,” Ultrason. Imaging, 2018.

[64] P. M. Gammell, “Improved ultrasonic detection using the analytic signal magnitude,” Ultrasonics, vol. 19, no. 2, pp. 73–76, 1981.

[65] N. A. Azman and S. B. Abd Hamid, “Determining the Time of Flight and Speed of Sound on Different types of Edible Oil,” in IOP Conference Series: Materials Science and Engineering, 2017.

[66] J. C. Bamber and C. R. Hill, “Ultrasonic attenuation and propagation speed in mammalian tissues as a function of temperature,” Ultrasound Med. Biol., 1979.

[67] J. P. Thirion, “Image matching as a diffusion process: An analogy with Maxwell’s demons,” Med. Image Anal., 1998.

[68] D. P. Kingma and J. L. Ba, “Adam: A method for stochastic gradient descent,” ICLR Int. Conf. Learn. Represent., 2015.

