## Supplementary material for "Multiplicative frequency and angular speckle reduction in ultrasound imaging": http://www.stanford.edu/group/chugroup/

*1.1. Experimental setup for phantom imaging and the alignment of the phantom images*

Figure S1A shows a schematic of the experimental setup for combined frequency and angle compounding. Figure S1B shows the frequency compounded speckle image of the phantom at 9 different angles. They correspond to the speckle images shown in Fig. 2 in the main text, but with an extended field of view. The images show the corners of the hyperechoic region. The blue crosses are plotted at the same positions in all images. The center of rotation of the digital image rotation is adjusted so that the corners of the hyperechoic region in all the images are aligned, as indicated by the blue crosses. This process aligns the center of rotation of the digital images with that of the sample as it was rotated in the experiment.

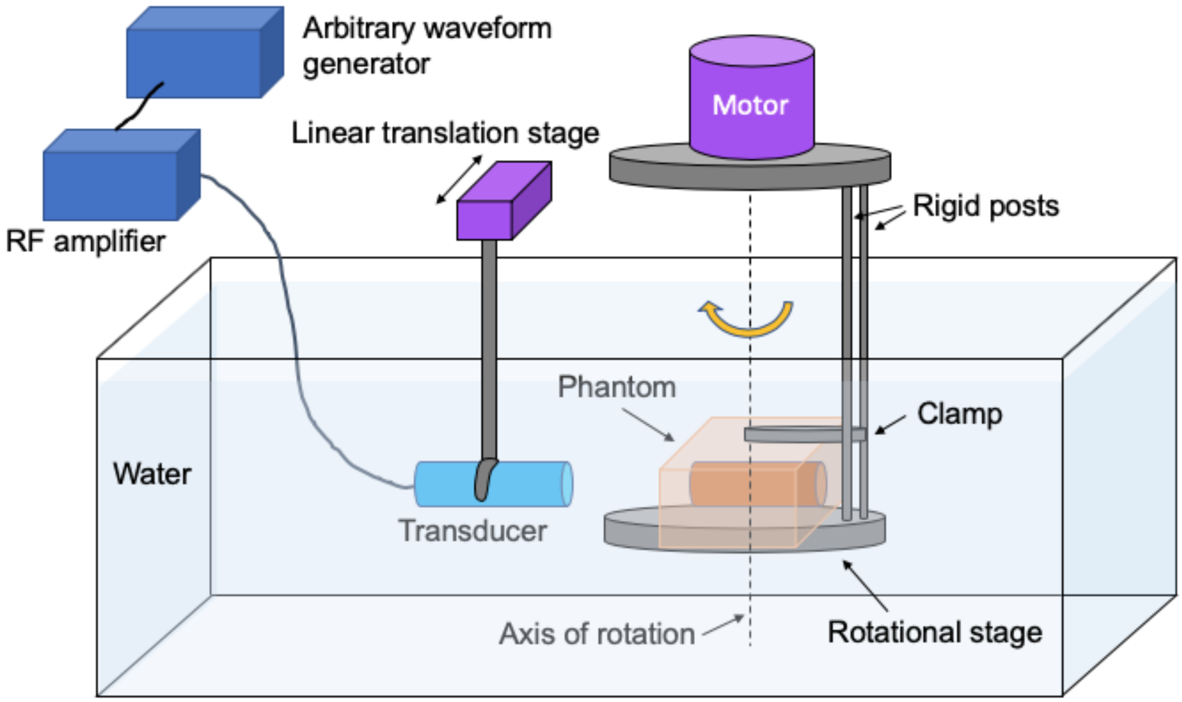


**Fig. S1A**. Experimental setup for phantom imaging. The phantom (shown in orange) is fixed on a rotational stage that is mounted to a motor, which rotates the phantom to a number of orientations with the rotation axis in the vertical direction. The transducer is oriented such that the transmitted sound pulse propagates in the horizontal direction. The linear translation stage moves the transducer perpendicular to the propagation direction of sound. The transducer is driven by amplified electrical waveforms produced by an arbitrary waveform generator.

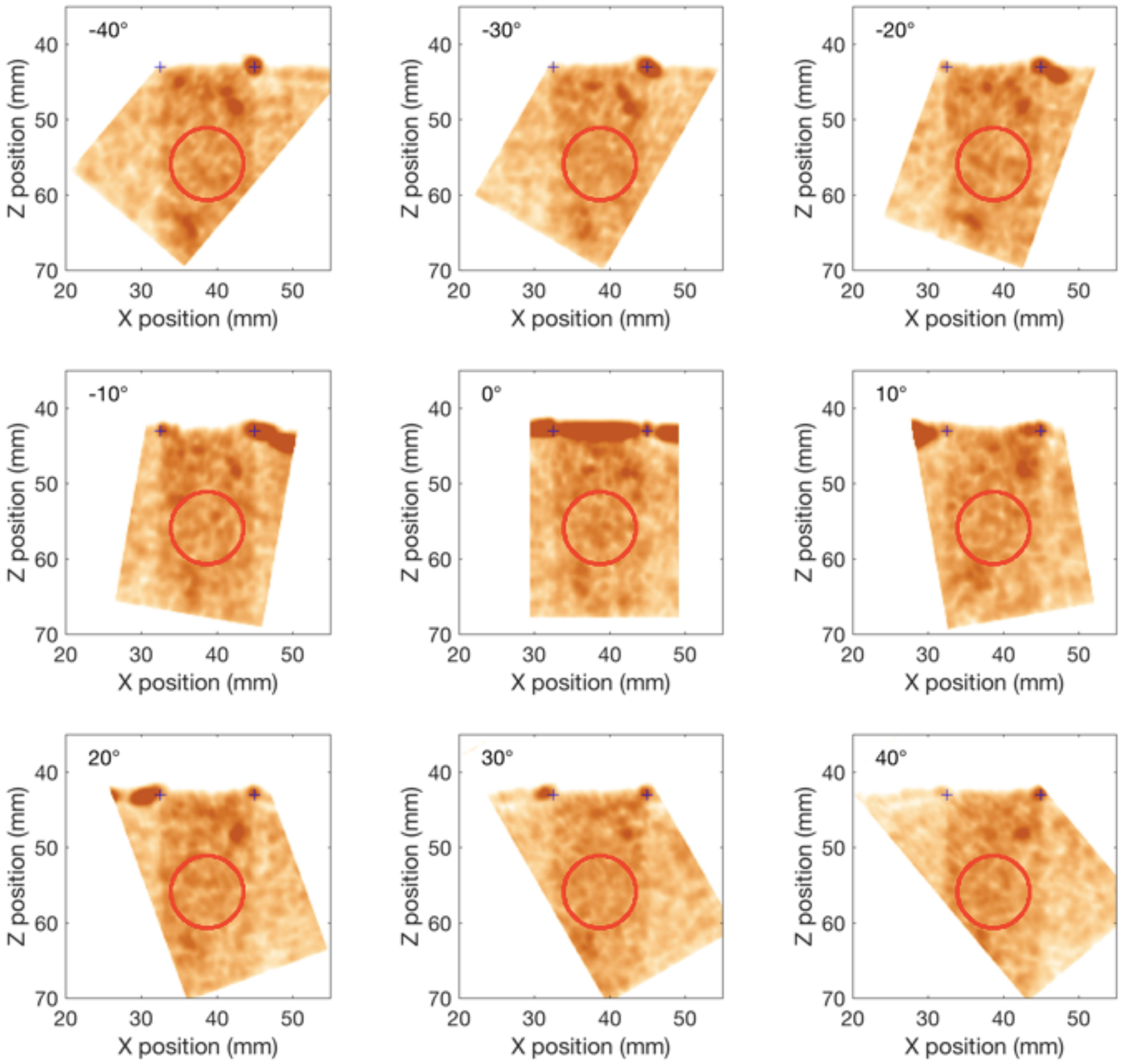


**Fig. S1B.** Frequency compounded images of an agarose phantom that is dispersed with corn starch particles. The center hyperechoic region has 3× higher corn starch concentration than the rest. Two blue crosses in each sub-figure are located at the same spatial coordinates in the figures ($x_1 = 32.5$, $x_2 = 45$, $z_1 = z_2 = 43$). The red circle indicates the region in which the speckle reduction is evaluated.

*1.2. Rigid alignment of the wrist images*

The frequency compounded images taken from different angles are rotated by the angle of incidence determined by the robot arm. Fig. S2 shows the compounded image after only applying the rigid transform. Misalignment can be seen near the skin on the right side of the image. This misalignment is due to the contact of the tissue by the ultrasound probe. Different regions in the compounded image are averaged from different numbers of images (Fig. S3).

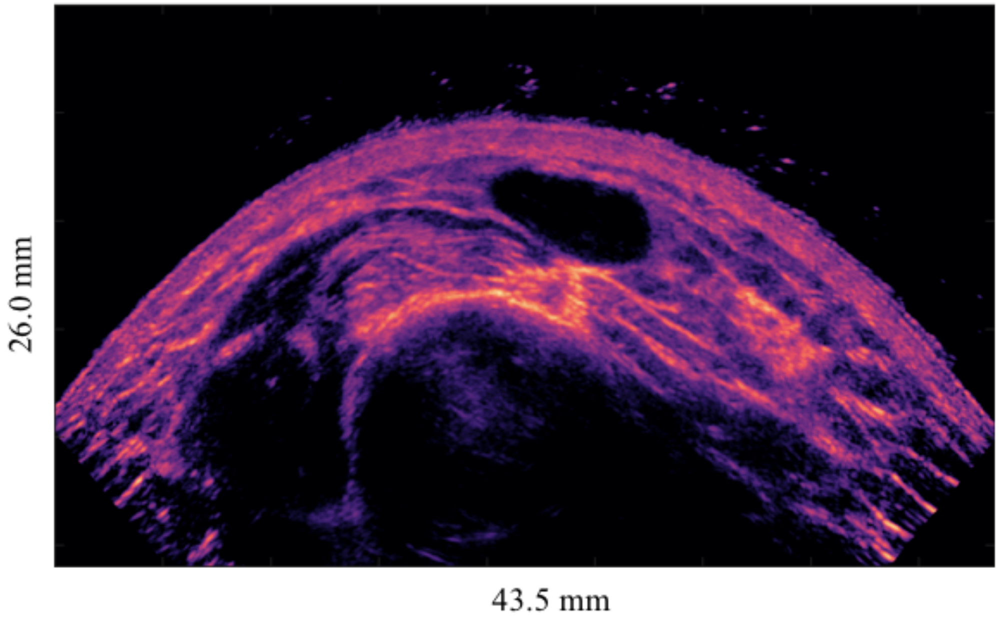


**Fig. S2**. Frequency and angle compounded image of the wrist. The rotated images are aligned using rigid rotation and translation matching.

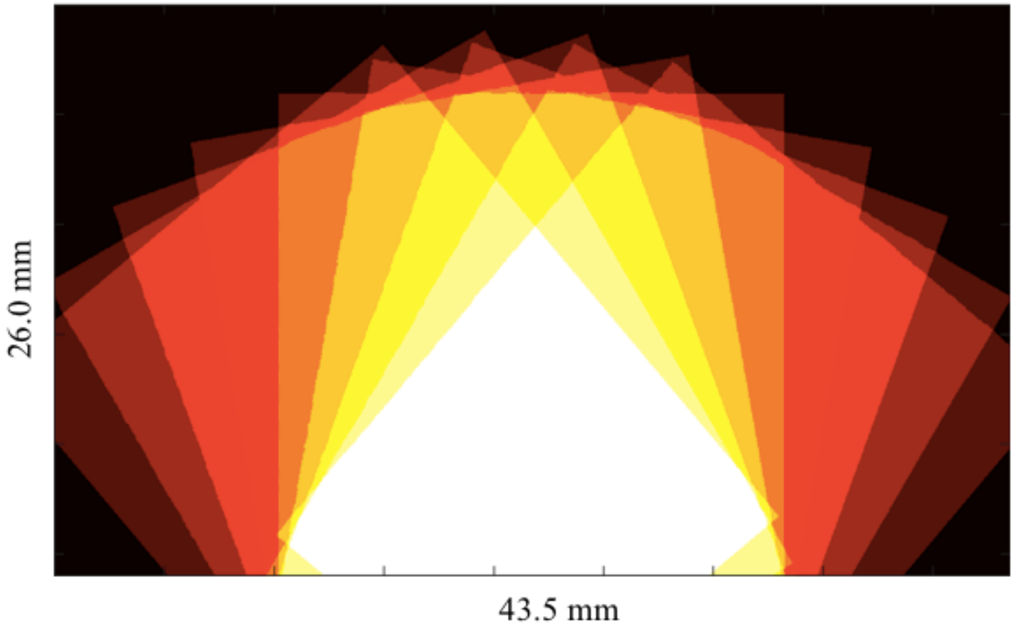


**Fig. S3.** False color plot of the degree of angle compounding. The color indicates the degree of image overlap ranging from 1 – 9 angles with increasing brightness.

*1.3. Trade-off between axial resolution and speckle reduction*

An optimized trade-off between axial resolution and speckle reduction can be obtained with Gaussian pulses. [1] Fig. S4 shows the frequency and angle compounded images with different bandwidths for frequency compounding. As the bandwidth increases, the axial resolution is improved, but the speckle reduction becomes less effective. The estimated total speckle reduction is 6.8×, 5.6×, 3.9×, 3.0× for the 1$\sigma$ bandwidths of 0.8, 1.2, 2.5, and 5.0 MHz, respectively.

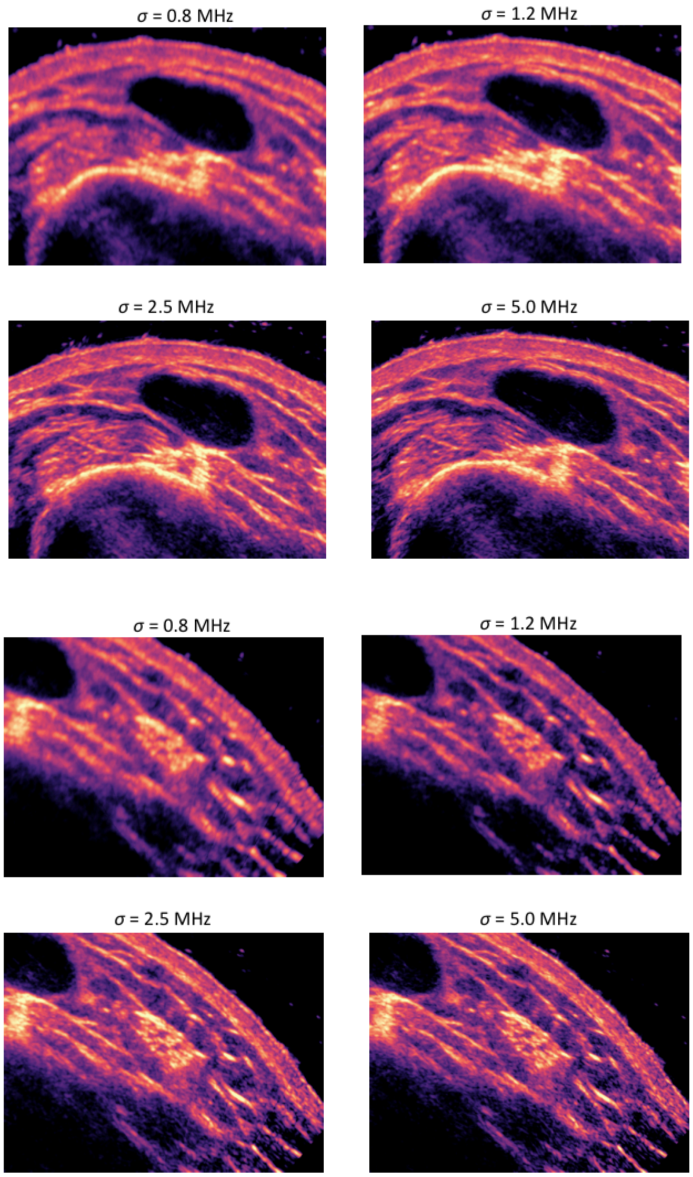


**Fig. S4.** False color images a human wrist compounded at 6 frequencies, 6.0, 8.0, …, 16.0 MHz and 9 angles −40, −30, …, 40°. Each panel corresponds to different widths used in frequency compounding. The top four panels and the bottom four panels show different regions of the wrist.

*2.1. Definitions for image and image transform*

"Image" and "image transform" are defined as the following. An "image" $I(x_i, y_i)$ is defined as a discretely sampled function on a coordinate space $R^2 = \{x_i, y_i\} \to R^n$, where the set $\{x_i, y_i\}$ are the coordinates of an array of pixels representing the image, and $R^n$are the $n$ "color" values for each pixel $x_j, y_j$. In the case of ultrasound imaging, the image is monochromatic and $n = 1$. Two different images then represent two maps on two different coordinate spaces. The "image transform" that allows the registration of two images is defined as the coordinate transformation $T(x, y): R^2 \to R^2$. In the case of a rigid translation, $T(x, y): (x, y) \to (x + \delta x, y + \delta y)$. The transform $T$ is used to synthesize a new image $I'((x_i, y_i) = I\big(T(x_i, y_i)\big) = I(x_i', y_i')$. For the rigid translation given above, $I' = I(x + \delta x, y + \delta y)$.

The goal of an image registration algorithm is to find a transformation $T$ that maps points in the coordinate space from one image onto the coordinate space of another image. One of the images is selected to be the "stationary" image $S$ defined in its coordinate system. The other image is designated as $M$, the "moving" image. The transformation $T$ that shifts the coordinates of $S$, $(x_i, y_i)$ to the coordinates of $M$, $(x_i', y_i')$. If there is a perfect correlation and registration between the two images, $S(x, y) = M(T(x, y))$. For convenience, we define $M' \equiv M(T(x, y))$. The non-rigid image registration problem is illustrated in Fig. S5 and photos of a toucan bird are used. While the current work is concerned with monochrome images, colors in the examples are kept and it is assumed that the corresponding grayscale values are used for image matching. Note that margins with zero-pixel values have been appended to the images to allow for distortions that are outside the original image boundaries. The transform $T$ in this example is parameterized by a set of displacement vectors $\{\boldsymbol{a}_i\}$ which map each image pixel $\boldsymbol{p'} = (x, y)$ in the image $M'$ to a corresponding point in $M$. In other words, the $\{\boldsymbol{a}_i\}$ represent a discretized $T$ which constructs an image by sampling $M$ such that the intensity value at $\boldsymbol{p'}$ in $M'$ is given by the value at $\boldsymbol{p}$ in M with $\boldsymbol{p} = \boldsymbol{p'} + \boldsymbol{a_i}$. (Fig. S6) Fig. S7 shows the spatial distribution of a subset of $\{\boldsymbol{a}_i\}$ and the deformed image $M'$.

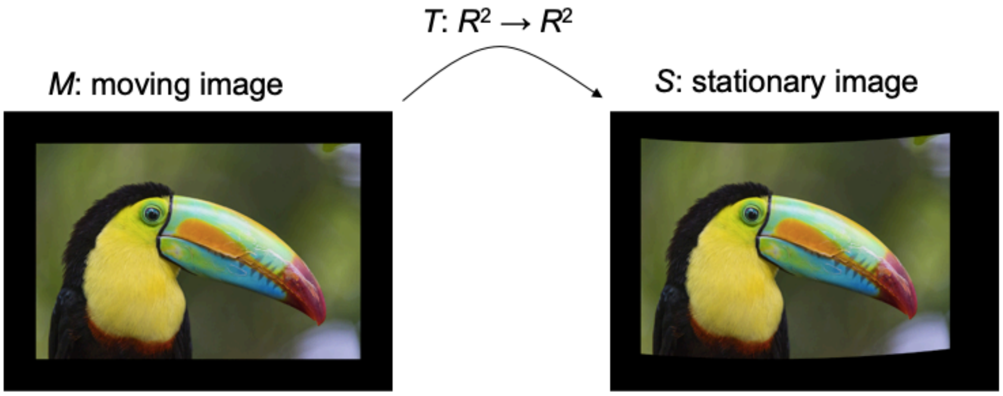


**Fig. S5.** The problem of nonrigid image registration concerns the transformation of a moving image *M* to match a stationary image *S*.

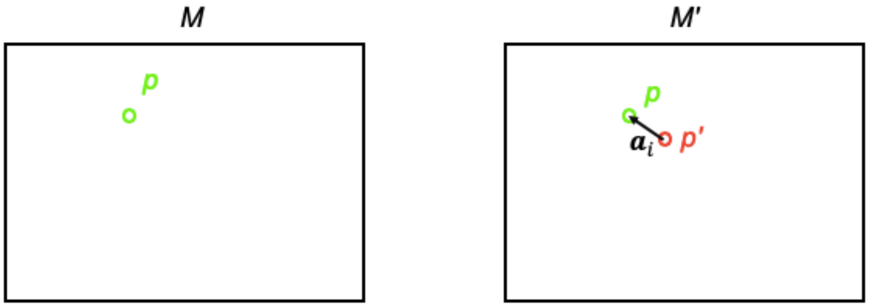


**Fig. S6.** A displacement vector $\boldsymbol{a}_i$ starts from $\boldsymbol{p}'$ in $M'$. The image value at $\boldsymbol{p}'$ is given by the image value at $\boldsymbol{p}$ in $M$.

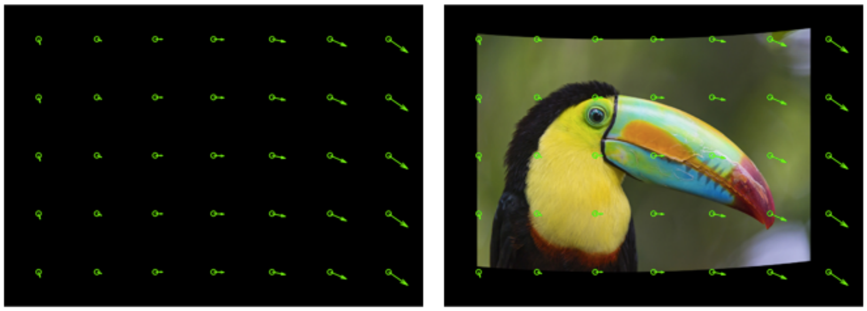

**Fig. S7.** A new moving image (right) is obtained from the displacement vectors $\boldsymbol{a}_i$ (left).

*2.2. Details of the image matching algorithms*

Two different iterative registration algorithms are explored to correct for nonrigid distortions. The first one is based on the "demons" algorithm [2] implemented in the "`imregdemons`" function in MATLAB. The second algorithm is based on stochastic gradient descent using the ADAM optimizer [3] and is implemented in python. The main differences of the two algorithms are in the methods the update displacement vectors and on the constraints used limit the amount of deformation, also referred to as "regularization" that imposes an Occam razor principle to minimize overfitting.

In the demons algorithm, a lattice of vectors on the grid points (each pixel) in $M$ parametrize the transform $T$. The set of individual pixel displacement vectors is denoted as $\{\boldsymbol{a}_i\}$. In each iteration, the $\boldsymbol{a}_i$'s are updated with $d\boldsymbol{a}_i$ using the optical flow equation [2]:

$$d\boldsymbol{a}_i \cdot (\boldsymbol{\nabla} s)_i = (s - m')_i, \quad (1)$$

$(\boldsymbol{\nabla} s)_i$ is the gradient of the image values $s_i$ at pixel *i*. The geometric interpretation of this equation is based on the linear approximation of the local image intensity in *S*, such that the update displacement vector $d\boldsymbol{a}_i$ will match the difference between $s$ and $m'$. The updated displacement vectors are smoothed by a Gaussian spatial filter, which has a $1\sigma$ width of 3 pixels, roughly corresponding to the image voxel size. The smoothing process regularizes the displacement field against variations in the displacement field at high spatial frequencies compared to the Gaussian width. The cycle is repeated for predetermined number of steps and the algorithm obtains the image $M'$ using the updated $\boldsymbol{a}_i$.

The optical flow equation (Eq. 1 in main text) plays a central role in the demons algorithm and can be understood intuitively from a geometric representation of the pixel values *s* in the 3D coordinate. (Fig. S8) The z-axis represents the value of *s* and the x- and y-axes represent the image pixel coordinates. $\boldsymbol{\nabla} s$ is the gradient of *s* in the x-y plane. The vector $(\boldsymbol{\nabla} s/|\boldsymbol{\nabla} s|, |\boldsymbol{\nabla} s|)$ then represents the slope of the *s*-surface at the point $\boldsymbol{p}'$. The goal of image matching is to find the vector in the *x-y* plane that matches $m'$ with *s*. To the first order approximation, this vector is obtained by extending the slope vector to intersect with the $s = m'$ plane (light blue plane in Fig. S8) at a point $\boldsymbol{q}'$. $\boldsymbol{q}' - \boldsymbol{p}'$ is then the required shift of the point $\boldsymbol{p}'$ in the *x-y* plane such that the *s*-surface touches the $s = m'$ plane to the linear approximation. This geometric consideration gives rise to the 'optical flow equation', which in this case is $(\boldsymbol{q}' - \boldsymbol{p}') \cdot \boldsymbol{\nabla} s = m' - s$. To apply the optical flow equation for updating the displacement vector, it should be taken into consideration that the vector $\boldsymbol{a}_i$ starts from $\boldsymbol{p}'$, instead of ending at $\boldsymbol{p}'$, hence the required update vector is $d\boldsymbol{a}_i = -(\boldsymbol{q}' - \boldsymbol{p}')$ and the desired optical flow equation (Eq. S1) is obtained. To invert the optical flow equation and obtain $d\boldsymbol{a}_i$, $\boldsymbol{\nabla} s$ is shifted to the right-hand side and a small regularization value is added to avoid the problem of zero denominator. [2]

In the gradient descent algorithm, a grid of 10×10 points $\{\boldsymbol{b}_i\}$ are selected to parametrize the displacement vectors $\{\boldsymbol{a}_i\}$. The updates to the $\boldsymbol{b}_i$'s are computed by a loss function *L*

$$L = L_{fit} + L_{reg}. \quad (2)$$

As an example, $L_{fit} = \sum(m' - s)^2$ uses a least squares criterion for the goodness of the registration. The second term, $L_{reg}$ imposes a regularization penalty to overfitting the distortions. The update vector $d\boldsymbol{b}_i$, which denotes the change in $\boldsymbol{b}_i$, is computed from the gradient of the loss function using the ADAM (adaptive moment estimation) optimizer [3]. To form the updated image, the $\boldsymbol{a}_i$'s are obtained by interpolating the values of the $\boldsymbol{b}_i$'s.

TABLE I

NON-RIGID IMAGE MATCHING ALGORITHMS.

| Demons algorithm | Gradient descent algorithm |
|---|---|
| The displacement vectors are initialized to be $\boldsymbol{a}_i = \vec{\boldsymbol{0}}$ for each pixel in *M'*. $\boldsymbol{\nabla} s$ is calculated with respect to the displacement vector $\boldsymbol{a}_i$ | A group of 10 by 10 grid points $\boldsymbol{b}_i$ are selected to parametrize and assigned with displacement vectors $\boldsymbol{b}_i$ . |
| Compute updated $a_i$'s by computing $d\boldsymbol{a}_i$ using optical flow (Eq. S1).<br>Smooth $\boldsymbol{a}_i$'s with a Gaussian spatial filter.<br>Obtain *M'* using updated $\boldsymbol{a}_i$ | Calculate the gradient of the loss function with respect to $\boldsymbol{b}_i$, and compute the update vector $d\boldsymbol{b}_i$.<br>Interpolate $\boldsymbol{b}_i$ to obtain the displacement vectors $\boldsymbol{a}_i$.<br>Obtain *M'* using updated $\boldsymbol{a}_i$ |

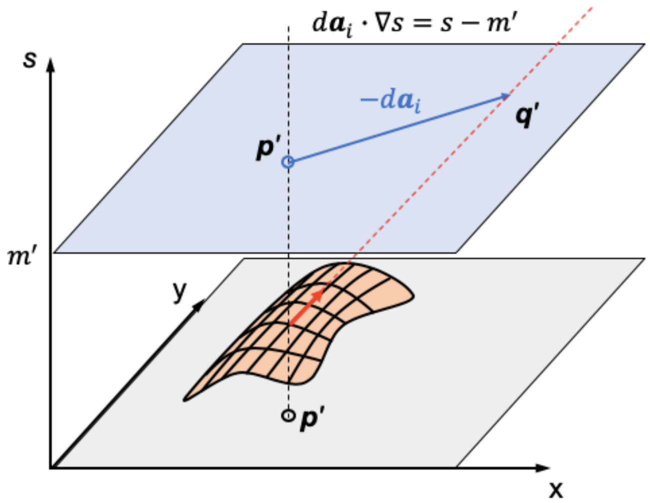


**Fig. S8.** The update vector $d\boldsymbol{a}_i$ is determined from the optical flow equation, which can be understood geometrically. The orange surface represents the pixel values of the stationary image S. The vector $(\nabla s/|\nabla s|, |\nabla s|)$ is shown in red. Its projection onto the x-y plane is along the gradient vector $\nabla s$. The surface corresponding to $s = m'$ is shown in blue. The extension of the tangent vector intersects with the $m'$ plane at point $\boldsymbol{q}'$. $\boldsymbol{q}' - \boldsymbol{p}'$ is thus the vector that matches $s$ and $m'$ to the linear approximation and the corresponding expression is $(\boldsymbol{q}' - \boldsymbol{p}') \cdot \nabla s = m' - s$.

To improve speed, both algorithms operate on down-sampled images. For the demons algorithm, 2048 and 128 iterations were run at 4× and 2× down-sampling, while the gradient descent algorithm ran 60, 30, 30 iterations at 8×, 4× and 2× down-sampling, respectively. The two algorithms are summarized in Table 1 in the main text. Good correction is achieved by both algorithms at a rate of a few seconds per image on a personal computer. It is expected that this process could be sped up significantly by using graphics cards and may be further improved by implementing in a faster language (e.g. C++) as compared to Python and MATLAB. Our results indicate that a fully optimized system will likely be able to operate in real time on currently available hardware.

*2.3. Regularization in the gradient descent algorithm*

The deformation field in the gradient-based model was parameterized by an evenly spaced grid of nodes. The displacement was interpolated using b-splines. Kybic, *et al.* [4] argue that B-splines are a suitable basis for non-rigid image registration modeling based on their expressiveness and the relative computational savings compared to other distortion field bases.

The deformation is regularized by including an additional loss term which models the severity of the distortion. The simplest version of this constraint is an energy defined by the interaction of nearest neighbor nodes. Explicitly, for any pair of nodes a penalty of the form below is computed:

$$E(x, y) = k((\mathrm{x} - \mathrm{x}_0)^2 + (\mathrm{y} - \mathrm{y}_0)^2)$$

where x and y refer to the displacement of the node relative to its starting position, while $x_0$ and $y_0$ refer to the average displacement of the surrounding neighbor nodes. Explicitly, when the displacement of a node is equal to the "equilibrium" position of the mesh at that point, the associated penalty is 0. As the node moves away from this "equilibrium" it gets assigned a penalty equivalent to that of a spring connecting it to the equilibrium point.

At the boundary of the field, the mean displacement is computed based on the adjacent nodes and does not consider the "missing" nodes (corresponding to points outside the field). Note that our model does not constrain the magnitude of the displacement field at any given location to avoid the limitations of physical elastic models that sometimes constrain the absolute value of the distortions [5].

The nearest neighbor model is not effective at constraining long (spatial) wavelength oscillations. This is evident since all information about long wavelength information has to be implicitly transmitted through pairs of neighbor interactions. In order to include long range interactions, (i.e. computing an additional loss for displacement relative to nodes beyond the nearest neighbors), a weighted average is computed for the displacements (for each x and y displacement) of the N spatially nearest neighbors weighted by a 2D hamming window.

*2.4. Convergence of the demons algorithm*

Convergence in the matching of two images using the demons algorithm is quantified by the maximum and average distortion vector length. The full resolution for the wrist images is 40 $\mu m$/pixel. At 4× lower image resolution, the rate of increase in both the maximum and the average distortion vector lengths (such as shown in Fig. S9) decreases with increasing iteration number (Top row of Fig. S9). 2048 iterations were used as a stopping point when the maximum distortion vector length is more than 90% of its length when 4× more iterations are taken. After 2048 iterations, it was found that the average distortion vector length is more than 70% of its length than if 4× more iterations. At 2× lower image resolution, an additional 128 iterations results in distortion vector lengths that are more than 95% of their values at 8× more iterations (Bottom row of Fig. S9). These results provide the rationale to use 2048 iterations at 4× reduced resolution and 128 iterations at 2× reduced resolution.

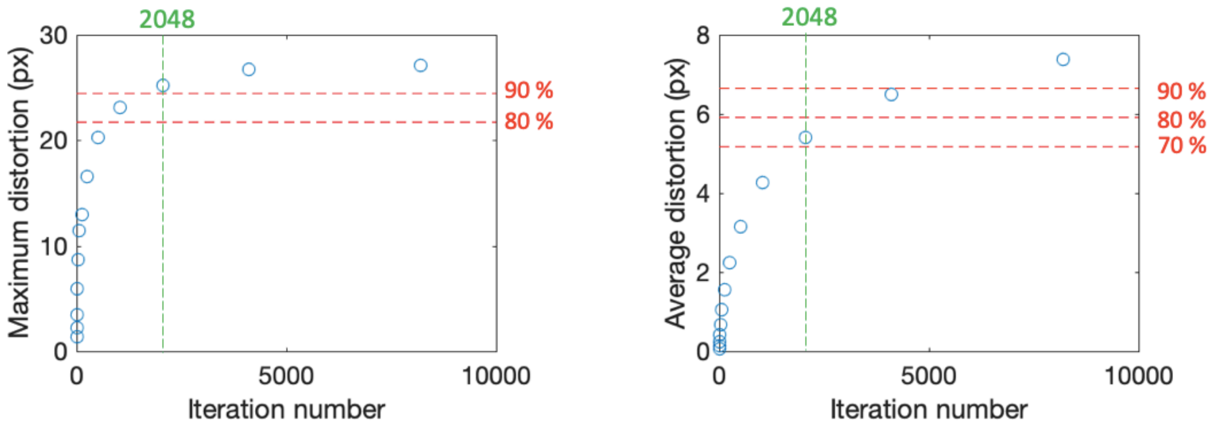

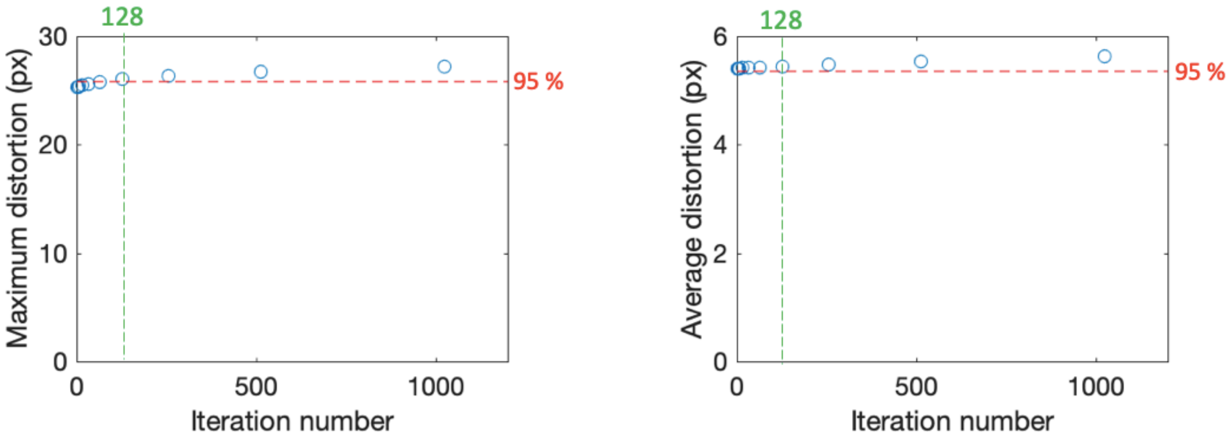


**Fig. S9.** The amount of maximum distortion and average distortion vector length as functions of iteration number for the matching of two wrist images at different angles. The top row shows the results with 4× reduced resolution. The bottom row shows the results with 2048 iterations at 4× reduced resolution and at 2× reduced resolution.

*2.6. Examples of under-constraint and over-constraint in the regularization schemes*

Fig. S10A shows the mismatch between the images after the nonrigid distortion with over-constrained "displacement" regularization, where the regularization term is amplified by 100× from the value used in the main text. Fig. S10C shows the unnatural "ripples" that are generated when the Gaussian smoothing is insufficient, corresponding to $1\sigma$ width of 1 pixel, as compared to 3 pixels in the adequately constrained case. Figs. S10B and S10D show the image matching results when the over-constrained and under-constrained regularization terms are corrected.

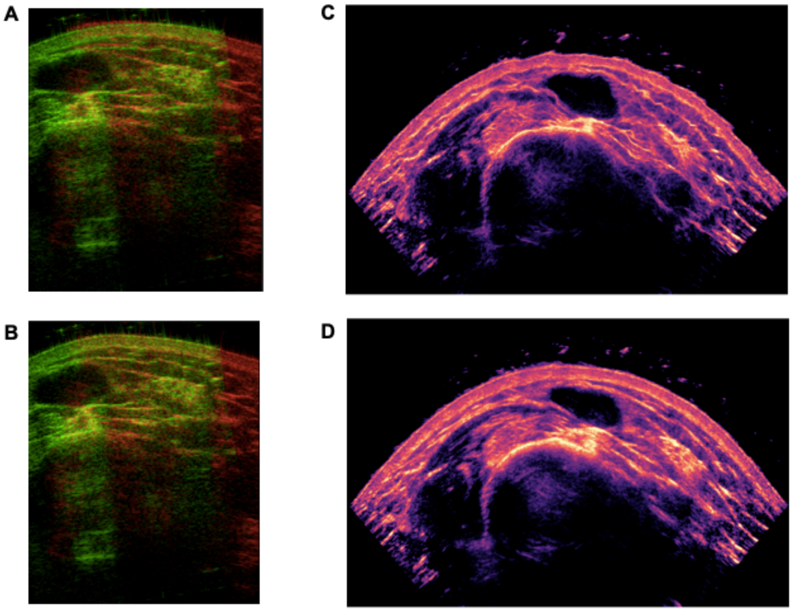


**Fig. S10.** False color wrist images with (**A**) over-constrained elastic loss function (where the red and green images do not overlap) and (**B**) adequate constraint obtained using the gradient descent algorithm. False color wrist images with (**C**) under-constrained and (**D**) adequate constraint obtained using the demons algorithm. Under-constrained fitting allows the algorithm to use excessive high spatial frequency corrections to make elastic distortions adjustments to a limited number of images. This is analogous to "over-fitting" a set of $N$ data points $\{x_i\}$ with a polynomial function $f(x)$ of order $n \geq N$.

*3. Regularization*

It is necessary to apply regularization to the image matching algorithms ensure robust and physically realistic solutions. The demons and gradient descent algorithms use, respectively, "velocity" and "displacement" regularizations. Velocity regularization constrains the magnitude of the *change* in the distortion map at each iteration step while displacement regularization constrains the magnitude of the distortion relative to the equilibrium state. Roughly speaking, these registration schemes can be compared to the flow of a viscous fluid and the deformation of an elastic solid. A viscous fluid can reach equilibrium in an arbitrarily complex shape but is constrained on how quickly it reaches this state by its viscosity. On the other hand, the elastic solid can be continuously deformed but is progressively more difficult to change as it is pushed beyond equilibrium.

An obvious downside of a displacement regularization is that even if a perfect registration state exists, the algorithm will not converge to it because there are two different loss terms that in general do not have the same minimum. The registration may get arbitrarily close when the regularization becomes arbitrarily weak, the relative magnitude of the registration loss to the over fitting penalty imposed by the regularization cost was determined empirically.

"Velocity" regularization has the drawback that the nonrigid distortion can be unnatural when the algorithm is under-constrained. Examples of under-constraint and over-constraint in the regularization schemes and how they are addressed are shown in Fig. S10.

*4. Side-by-side comparison of the registered wrist images by two algorithms*

Frequency and angle compounded wrist images are obtained by the demons and the gradient descent algorithms (Fig. S11). While there are fine differences in the resulting images, both algorithms demonstrate effective correction of the non-rigid distortions.

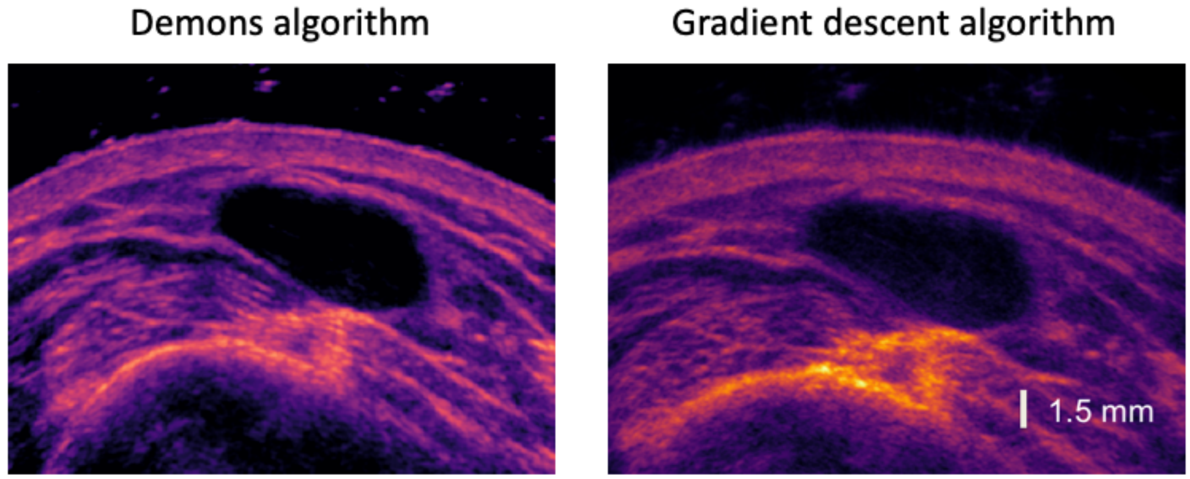


**Fig. S11**. Side-by-side comparison of frequency and angle compounded wrist images obtained by the demons and the gradient descent algorithms.

*5. An overall imaging framework*

The overall imaging framework is shown schematically in Fig. S12. This framework uses image processing to track a ROI to control an US device and registers and compounds images for speckle reduction. The framework can be broken down into a series of image processing and analysis operations which interface with US hardware to analyze and control the US.

A fast optical flow analysis of the continuous stream of US images captured by the US device enables both the ROI tracking and beam steering and image selection routines. An algorithm selects images for compounding from the continuous stream of US images which match a predefined criterion (e.g. a threshold for angular displacement from previous images). Attempting to register all images would be too computationally expensive and preclude real-time performance. This component allows the system to select images to maximize speckle reduction while remaining computationally tractable.

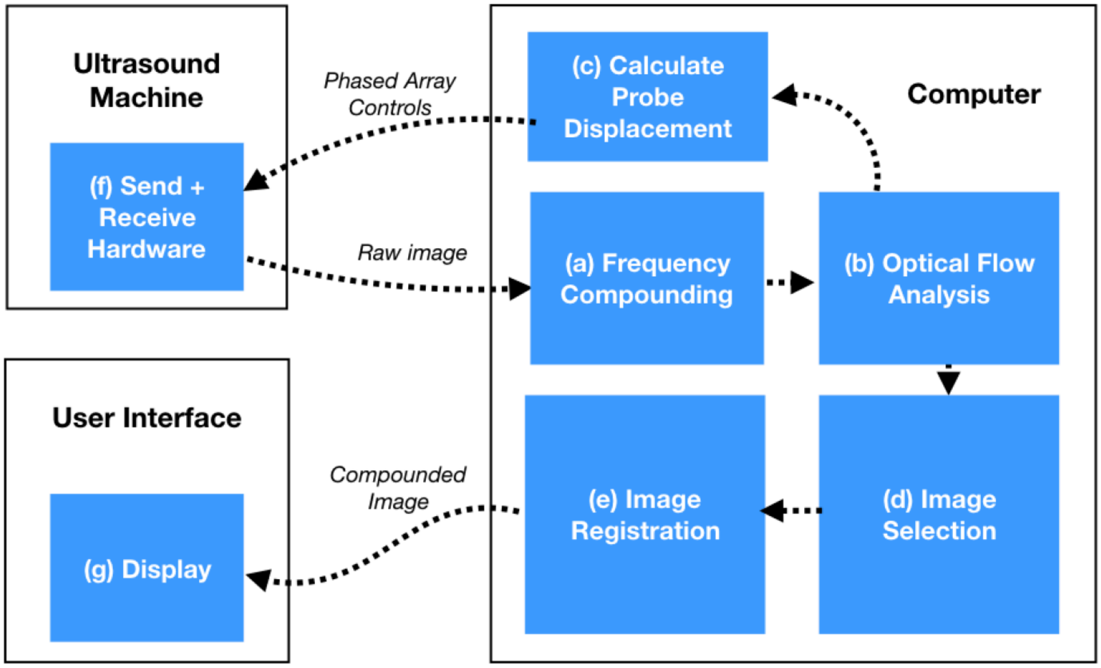


**Fig. S12.** Work-flow of image acquisition of a ROI. The ultrasound head is used to transmit and receive wideband ultrasound signals. Fast-Fourier transform, and Fourier-filter electronics can create frequency compounded images in real time. In Fig. 10 in the main text, the beam steering of the red image to overlap the ROI need not be precise. Instead of using the full number of pixels of the image, down-sampled image can be created by for example binning 4×4 pixels into a single pixel to form a lower resolution image that is used to calculate the phase delay need to center the ROI. Only a small fraction of the images collected in real time are used for image registration to create the final angle and frequency compounded image.

**References**

[1] Y. Li, Y. Winetraub, O. Liba, A. De La Zerda, and S. Chu, “Optimization of the trade-off between speckle reduction and axial resolution in frequency compounding,” *IEEE Trans.*

*Med. Imaging*, vol. 38, no. 1, 2019.
[2] J. P. Thirion, "Image matching as a diffusion process: An analogy with Maxwell's demons," *Med. Image Anal.*, 1998.
[3] D. P. Kingma and J. L. Ba, "Adam: A method for stochastic gradient descent," *ICLR Int. Conf. Learn. Represent.*, 2015.
[4] J. Kybic and M. Unser, "Fast parametric elastic image registration," *IEEE Trans. Image Process.*, 2003.
[5] G. E. Christensen, R. D. Rabbitt, and M. I. Miller, "3D brain mapping using a deformable neuroanatomy," *Phys. Med. Biol.*, 1994.
